# Enhancing the Understanding of Environmental Microbiomes through Topic Modeling: A Quantitative and Qualitative Analysis

**DOI:** 10.64898/2026.04.28.721390

**Authors:** Anna S. Kujat, Christiane Hassenrück, Stefan Lüdtke, Matthias Labrenz, Theodor Sperlea

## Abstract

**Background:** Understanding ecosystem dynamics is essential for assessing ecosystem health, yet remains challenging due to complex biotic and abiotic interactions. Microbial communities are valuable indicators of environmental change, but the high dimensionality of microbiome data requires advanced analytical methods. This study explores the use of topic modeling (TM), an unsupervised machine learning approach initially designed for text analysis, to analyze microbiome data from the dynamic Warnow Estuary on the southern Baltic Sea coast.

**Results:** We applied TM to estuarine microbiome data and compared its performance to traditional dimensionality reduction methods, Principal Component Analysis (PCA) and Principal Coordinate Analysis (PCoA). Quantitative results indicate that TM performs comparably to conventional approaches in preserving ecological and functional information, and in certain aspects even superior. In addition, we show qualitatively that NNMF, a TM method, captures latent patterns in the data providing an interpretable perspective on the microbiome. In this exploratory framework, NNMF suggested five distinct sub-communities within the estuary that appear to follow a seasonal succession influenced by freshwater inflow. These sub-communities were associated with specific ranges of salinity and temperature and showed distinct taxonomic profiles, with shared characteristics across the estuarine system.

**Conclusions:** Our findings suggest that TM is a useful tool for exploring complex environmental microbiome datasets, offering a complementary perspective that can provide additional ecological insights. TM’s ability to highlight coherent microbial community patterns indicates its promise for supporting environmental monitoring and informing targeted ecosystem management in dynamic habitats, though further studies are needed to fully assess its applicability.

## 1 Introduction

A comprehensive understanding of ecosystems and their dynamics is essential to accurately assess their functioning and their capacity to respond to and recover from disturbances. However, this understanding is challenging, as ecosystems are inherently complex, with their functioning driven by the interplay of biotic and abiotic components, variability across spatial and temporal scales, and feedback loops that can amplify or dampen changes (Anand et al. (2010); Riva et al. (2023)). A way to unravel ecosystems dynamics lies in understanding their biotic communities, as they are integral components of every ecosystem. Because biota respond distinctly to environmental changes, analyzing their spatial and temporal dynamics can provide insights into underlying ecological processes. For instance, bioindicator-based approaches have been used to assess water quality and trace metal concentrations in aquatic systems (Miller et al. (2007); Barinova (2017); Barinova et al. (2020); Dokulil (2003)). Among these bioindicators, microbial communities—fundamental to all ecosystems—are particularly well-suited for environmental monitoring due to their ubiquity, short generation times, and sensitivity to environmental changes (Janßen et al. (2021), Li et al. (2020a), Sagova-Mareckova et al. (2021)).

The establishment of environmental DNA (eDNA) protocols combined with culture-independent next-generation sequencing (NGS) has revolutionized the study of microbial communities (Sagova-Mareckova et al. (2021)). NGS enables the high-throughput simultaneous analysis of thousands of microbial features per sample, producing microbiome tables (*n* × *p*) where each cell represents the relative abundance of a specific organism (*p*) in a given sample (*n*) (Hugerth and Andersson (2017)).

However, NGS-based microbiome datasets pose significant analytical challenges. Beyond the inherent non-linear relationships and the compositional nature of the data, their high dimensionality adds significant complexity (Tsilimigras and Fodor (2016); Verleysen and François (2005)). In high-dimensional data, the number of detected organisms often exceeds the number of samples (*n* ≪ *p*), giving rise to the so-called “curse of dimensionality”. This phenomenon increases data sparsity and computational costs due to the large number of features, while a small sample size limits generalization to other datasets (Medina et al. (2022)). As a consequence, traditional statistical methods like linear models are prone to overfitting and often fail to capture the non-linear relationships essential for understanding microbial systems (Verleysen and François (2005), Kiselev et al. (2019)).

Thus, advanced methods are required to reduce the dimensionality of NGS-based microbiome data while preserving key ecological patterns. Principal Component Analysis (PCA) is a widely used dimensionality reduction method (DRM) in ecological studies, summarizing high-dimensional data into a set of *k* components, where *k* ≪ *p* (James and Mcculloch (1990)). However, PCA often fails to capture the non-linear relationships that are fundamental to understanding complex ecological systems (Verleysen and François (2005); Kiselev et al. (2019); Sperlea et al. (2021)). Principal Coordinates Analysis (PCoA), by contrast, does not rely on linear assumptions and is therefore commonly used as a dimensionality reduction method in microbiome research. Unlike PCA, PCoA operates on a distance matrix—typically based on dissimilarity measures such as Bray-Curtis—and seeks to preserve these pairwise distances when projecting the data into a lower-dimensional space (Paliy and Shankar (2016); Medina et al. (2022); Daniels et al. (2014)). However, despite its advantages, PCoA does not provide insight into the contribution of individual features (e.g., amplicon sequence variants, ASVs) to the observed patterns.

More recently, machine learning (ML)-based dimensionality reduction methods have been increasingly adopted in ecological research (Medina et al. (2022); Pichler and Hartig (2023); Cordier et al. (2019)). One such approach is topic modeling (TM)—a class of unsupervised ML algorithms originally established in natural language processing—which makes no explicit linear assumptions and simultaneously provides interpretable feature distributions. Unlike PCA or PCoA, TM generates low-dimensional representations of a dataset’s feature space by clustering co-occurring elements (e.g., words in documents) into a smaller number of latent groups (*k* ≪ *p*), referred to as *topics*, each representing a coherent structure shared across the data entities (*n*). This principle extends to ecological data, where co-occurring organisms may indicate shared ecological roles. Latent Dirichlet Allocation (LDA), a probabilistic TM method, assumes that documents are mixtures of topics, where words are assigned to topics probabilistically (Blei et al. (2001)). LDA has already been applied in ecology to classify forest types, link animal populations to climate events, and analyze plant microbiomes in relation to host stress levels (Valle et al. (2014); Christensen et al. (2018); Hosoda et al. (2020); Kim et al. (2023)).

An alternative, non-probabilistic TM approach is Non-Negative Matrix Factorization (NNMF). First introduced in image processing (Lee and Seung (1999)), NNMF has since gained popularity in text analysis, establishing itself as a key TM approach (Vayansky and Kumar (2020); Xu et al. (2003)). Its non-negativity constraint aligns well with ecological data, where abundance measures are inherently non-negative. NNMF has been used to uncover molecular patterns in cancer genomes and microbial features in mammalian stool samples, providing valuable insights into biological processes (Brunet et al. (2004); Cai et al. (2017)).

Despite initial applications in biological and ecological research, TM remains under-utilized in environmental microbiome research, especially in highly dynamic ecosystems. Estuaries—transitional zones between freshwater and marine environments—are such complex habitats with steep salinity gradients and high spatiotemporal variability. While providing essential ecosystem services, including high primary production and natural buffering capacity, estuaries are increasingly affected by human activities (Dafforn et al. (2012); Pinckney et al. (2001); Day et al. (2012)). To date, studies on microbial diversity in estuarine systems have primarily focused on specific time points or seasons (Crump et al. (2004), Bouvier and Giorgio (2002), Zhang et al. (2006), Cottrell and David (2003)) and often concentrated on variations along salinity gradients—a key factor influencing aquatic microbiomes (Herlemann et al. (2011)).

To demonstrate the utility of TM in a highly dynamic setting, we applied it to prokaryotic microbiome data from the Warnow Estuary and the adjacent southwestern Baltic Sea coast. The Warnow Estuary is a relatively short (14 km), eutrophic system with circulation dynamics primarily driven by wind rather than tidal forces (Doherty et al. (2017), Campbell and Kirchman (2013), Lange et al. (2020)). This leads to rapid and strong fluctuations in water conditions, preventing the establishment of a stable salinity gradient. Beyond its variable salinity, the estuary is also influenced by multiple anthropogenic pressures, including urban wastewater discharge, industrial activity, and tourism (Lange et al. (2020)). Thus, the Warnow Estuary serves as an ideal difficult test case for exploring microbial dynamics in a highly dynamic ecosystem supported by TM approaches. Our microbiome dataset covers 14 locations along the estuary and the adjacent habitats and includes a full year of twice-weekly sampling, providing high spatial and temporal resolution of bacterial community dynamics.

To assess TM’s effectiveness, we employed a two-stage evaluation framework (Figure 1). First, we conducted a quantitative assessment to determine the optimal DRM for this environmental microbiome dataset by comparing TM with the conventional PCA and PCoA approaches. This framework accounts for ecological and functional properties of microbial communities, evaluates different data preprocessing strategies, and examines how the number of clusters (*k*) influences model performance. In the second stage, we selected a solidly performing TM approach based on the quantitative evaluation step to conduct an exploratory qualitative analysis. Using this approach, we revealed spatially distinct microbial sub-communities and additionally seasonal succession patterns, abiotic preferences, and taxonomic structures unique to these sub-communities, offering novel insights into microbial ecology in a dynamic system. Overall, this study demonstrates the potential of TM as a complementary tool to conventional approaches, enhancing our understanding of microbial dynamics in ecologically complex ecosystems, thereby providing a foundation for ecosystems state assessments, using the Warnow Estuary as a case study.

**Fig. 1.**
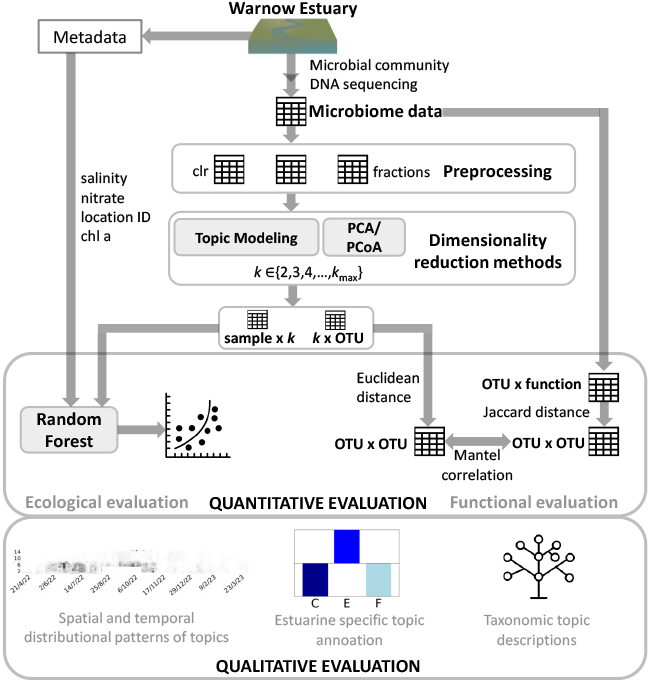
Graphical summary of the topic modeling approach for environmental microbiome data from the Warnow Estuary.

## 2 Methods

### 2.1 Sample Collection and Metadata Acquisition

The samples on which the microbiome dataset is based were collected between April 2022 and April 2023 as part of the OTC-genomics project (Sperlea et al. (2025)). A total of 1551 water samples were gathered through twice-a-week sampling, conducted consistently on Mondays and Thursdays, starting three hours after sunrise. Sampling occurred at 14 fixed locations along the southern Baltic Sea coast and the Warnow Estuary (Figure 2). In addition to *in situ* measurements using a CTD probe, laboratory analyses were performed to quantify the concentrations of chlorophyll a and nutrient levels, including phosphate, nitrate, nitrite, and ammonium (Sperlea et al. (2025)). Additionally, we obtained freshwater inflow volumes into the Warnow Estuary from the Landesamt für Umwelt, Naturschutz und Geologie. This inflow data originate from a sampling site located downstream of the estuary on the Warnow River.

**Fig. 2.**
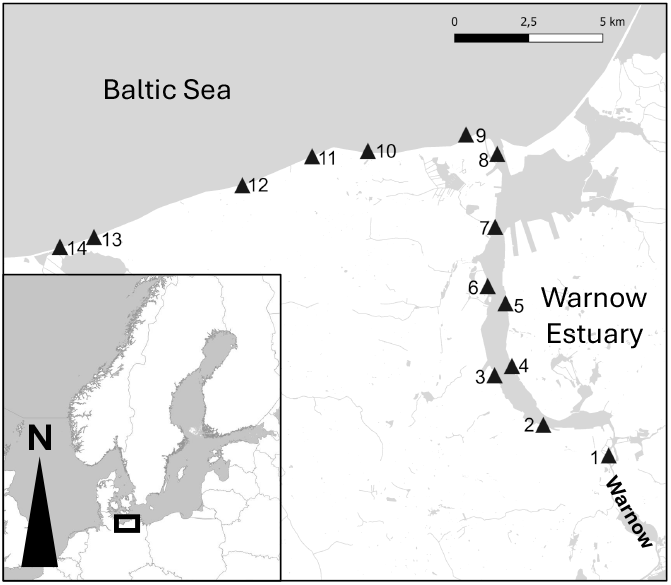
The sampling locations along the Warnow Estuary and the Baltic Sea coast.

### 2.2 DNA Extraction, Amplicon Sequencing and Raw Sequences Processing

Details on DNA extraction, amplicon sequencing, and raw sequence processing are provided in Sperlea et al. (2025). Importantly, the ASV table used in our analysis had already been filtered for rarity, excluding ASVs that did not occurred in at least 1% of samples with a relative abundance below 0.1%, resulting in a reduction of zero values by 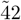 %.

### 2.3 Data Preprocession

NGS-based taxonomic ASV matrices contain compositional data, as they capture relative rather than absolute abundances. Since the total sequencing depth per sample is constrained and varies across runs, counts are only informative relative to each other. Moreover, due to incomplete sequencing of microbial communities and limitations of reference databases, absolute abundances cannot be reliably inferred (Tsilimigras and Fodor (2016)). Recent developments in introducing quantitative standards into microbiome sequencing workflows aim to overcome this limitation and may allow more accurate estimation of absolute microbial abundances in the future (Wang et al. (2025)). Here, to assess how the use of relative abundance data might affect TM analysis, we processed the microbiome dataset in three different ways. These included: (1) a centered log-ratio transformed matrix, which applies a log-ratio of each value relative to the geometric mean of all values in a sample, making the data suitable for compositional analysis (hereafter referred to as *clr* ). To handle zero values, we added a pseudocount of 0.5 prior to transformation; (2) a matrix representing the relative abundance of each ASV in relation to the total ASVs (hereafter referred to as *fractions*); and (3) using the original, unmodified matrix (hereafter referred to as *none*).

### 2.4 Dimensionality Reduction Methods

Here, the number of features, represented by ASVs, serves as the dimension to be reduced to a quantity *k* using DRM. The original microbiome dataset comprised 1551 samples × 1236 ASVs (Sperlea et al. (2025)), such that *k* is chosen to yield a substantial dimensionality reduction with respect to the full feature space. We compared TM, known for its application in the computational linguistic domain, with PCA as well as PCoA, two commonly used techniques in multivariate statistics for analyzing high dimensional ecological data.

#### 2.4.1 Topic Modeling

The TM models’ underlying concepts are transferable to our microbiome data because both ecological datasets and linguistic datasets can be represented in a similar mathematical form, a *n* × *p* matrix. In case of our microbiome dataset, *n* represents the samples, and *p* represents the ASVs. We applied the TM models LDA and NNMF. Probabilistic TM models, like LDA (Blei et al. (2001)), use probability distributions, with LDA defining topics as distributions over words. Translated to our microbiome dataset, this means that topics are represented as probability distributions over ASVs (*topic* × *ASV* ), and each sample is described as a probability distribution over topics (*sample* × *topic*). In contrast, non-probabilistic models such as NNMF rely on matrix decomposition techniques. NNMF, grounded in algebraic principles, enforces a non-negativity constraint, ensuring that each sample is represented as a non-negative combination of topics (*sample* × *topic*), and that each topic is composed of non-negative ASV contributions (*topic* × *ASV* ). As a result, both LDA and NNMF, with their different underlying principles, iteratively estimate two output matrices (*sample* × *topic* and *topic* × *ASV* ) using an optimization process, starting with random initial values, and refine these matrices until convergence is achieved. For simplicity, we refer to the ASV assemblages derived from TM topics as sub-communities. This terminology is used to highlight that these groups represent model-inferred assemblages of co-occurring taxa, rather than ecological communities in the strict sense. Throughout the manuscript, the term ‘sub-community’ therefore denotes topic-derived assemblages, i.e., sets of ASVs that tend to co-occur across samples as identified by the topic model.

Both TM implementations were from `sklearn.decomposition` of the `scikit-learn` package (version 1.3.2) (Pedregosa et al. (2011)). We used the models’ default settings: For LDA, we applied the `LatentDirichletAllocation` implementation with the number of components set to *k*, the number of total samples specified accordingly, and the random state fixed to 0. For NNMF, we used the `NMF` implementation with *k* components, initialized randomly, and again with the random state fixed to 0.

#### 2.4.2 Principal Component Analysis and Principal Coordinate Analysis

PCA as well as PCoA were performed to reduce the dimensionality of the microbiome data and to identify the main axes of variation across samples, enabling comparison with topics identified by TM. PCA produces both the coordinates of samples in the reduced space (*sample* × *component*) and the contributions of features (ASVs) to each component (*component* × *ASV* ). In contrast, PCoA provides only the sample coordinates in the reduced space (*sample* × *coordinate*). For PCA, we applied all three preprocessing approaches (*clr, fractions, none*) to allow a direct comparison with TM. The PCA implementation is from the `sklearn.decomposition` module with default settings in the `scikit-learn` package (version 1.3.2) was used (Pedregosa et al. (2011)). We used the `PCA` implementation with the number of components set to *k*. For PCoA, we first computed the Bray–Curtis dissimilarity matrix using `scipy.spatial.distance.pdist` and `squareform`, followed by applying multi-dimensional scaling (MDS) via the `MDS` class from `sklearn.manifold` (`scikit-learn` version 1.3.2) with *k* components, dissimilarity set to precomputed, and the metric option enabled. The resulting coordinates were obtained with the `fit_transform` function. Because the calculation of a Bray-Curtis dissimilarity matrix requires input data in a relative format, we used PCoA only with the *fractions*-transformed data.

### 2.5 Quantitative Evaluation

To quantitatively compare TM and PCA across different *k* settings and preprocessing strategies, we employed both an ecologically and a functionally motivated evaluation approach. These quantitative analyses provide an objective, reproducible basis to compare the performance of different dimensionality-reduction methods in capturing ecological and functional patterns in the microbiome.

#### 2.5.1 Ecological Evaluation

We employed Random Forest (RF), a supervised ensemble learning method, to assess the performance of various DRM approaches in an ecologically relevant context. Supervised machine learning models, like RF, identify patterns in data matrices alongside a target variable, build a model to capture these patterns, and predict the target variable for new, unseen data of the same type. In our setup, the abundance-derived data matrices (full ASV matrix or dimensionality-reduced representations) were used as predictors, while each environmental metadata variable was modeled separately as the target. This means that RF models were trained to predict environmental parameters from community composition.

For this, we generated *n* × *k* matrices using TM- or PCA/PCoA-based transformations, corresponding to different preprocessing methods and values for *k* that range between 11 and 191 in steps of 10 (*k* ∈ { 11, 21, …, 191 }). These matrices (*sample* × *topic, sample* × *PCA/PCoA component/coordinate*) were directly used as RF input features. No additional aggregation across samples was performed; each sample was represented by its vector of topic proportions (for TM) or component/- coordinate scores (for PCA/PCoA). The range of *k* ∈ { 11, 21, …, 191 } was selected arbitrarily in order to start with a significantly reduced dimension *k* and for assessing whether model performance peaked at specific values or approached saturation.

Because the data are time-resolved and exhibit seasonal trends, we did not perform a random train-test split. Instead, training and test sets were selected to respect the temporal order of the samples, with the training dataset comprising samples from April 2022 to January 2023, and testing conducted on samples collected from January 2023 to April 2023. This ensures that the model is evaluated on “future” data relative to the training set, providing a more realistic assessment of predictive performance across time (Regier et al. (2023); Boser (2024)).

As target variables, we included various metadata variables. For regression-based RF models, the continuous environmental variables salinity, chlorophyll a, water temperature and nutrient concentrations (nitrate, nitrite, ammonium and phosphate) were used, while for classification, the categorical variables location ID (1-14) and sampling week (1-52) served as targets.

To evaluate whether dimensionality reduction preserves ecologically relevant information, we compared the predictive performance of RF models trained on dimensionality reduced matrices against RF models trained on the full-dimensional *sample* × *ASV* matrix. The full ASV matrix represents an upper performance bound (since it retains all observed information), and thus served as a natural comparative baseline. Prior to this comparison, the full-dimensional matrix was preprocessed using one of three approaches—(*clr, fractions* or *none*)—and the best-performing preprocessing method was selected as the comparative baseline per target variable. This relative performance served as an indicator of which DRM approach preserved the most ecological information, encoded by the different environmental target variables. Importantly, the aim of this comparison was not to treat the full matrix as a reference point to be surpassed, but rather to assess how much predictive power is retained when using lower-dimensional DRM representations.

As performance measures, we initially used R^2^ for regression tasks and accuracy for classification tasks. For the more in-depth analysis of the numerical target variable chlorophyll a, we chose the mean absolute error (MAE) and the root mean squared error (RMSE) to supplement the R^2^, and calculated 95% confidence intervals using a bootstrap procedure with 1,000 iterations. We then performed an one-sample t-test on the bootstrapped R^2^ distributions of approaches, which performed better than the baseline to assess whether their performance differed significantly from the full microbiome baseline.

Target variables with a baseline predictive performance below 0.15 (R^2^, accuracy) were excluded from further analysis, as their limited predictability under the temporal split (likely due to strong seasonal variability within the single year of data) did not provide a meaningful basis for assessing DRM-based predictions. We used the RF evaluation implementation from scikit-learn (Version 1.3.2) (Pedregosa et al. (2011)) using the default model settings. For Random Forest classification, we applied the `RandomForestClassifier` with the random state fixed to 42. For Random Forest regression, we used the `RandomForestRegressor`, also with the random state set to 42.

#### 2.5.2 Functional Evaluation

To further evaluate the dimensionality reduction methods, we assessed how well TM and PCA preserve the functional properties of the microbiome after transformation. The underlying assumption of this analysis is that ASVs with similar functional roles should be grouped within the same topics, reflecting ecologically meaningful structure in the reduced feature space.

We used FAPROTAX, a database-driven tool for assigning prokaryotic taxa to metabolic functions (e.g., nitrification, fermentation), to create a functional profile for each ASV (Louca et al. (2016)). Importantly, these annotations reflect the genomic potential for metabolic functions based on taxonomic information and do not represent directly measured functional activity or gene expression. This functional assignment of the *sample* × *ASV* matrix resulted in a binary *ASV* × *function* matrix (0, 1: presence, absence of functions across ASVs).

DRM approaches generate either *topic* × *ASV* (TM) or *component* × *ASV* (PCA) matrices for each *k* (PCoA was excluded as it does not provide feature-level representations). In TM, these matrices assign ASVs to latent topics, while in PCA they represent ASV loadings on orthogonal components. Accordingly, the meaning of *k* differs: in TM it specifies the number of topics, whereas in PCA it denotes the number of retained components, with most variance usually explained by the first few. Despite these conceptual differences (topic structure vs. variance projection), both methods yield topic/component representations of the ASVs.

From these representations, we computed pairwise Euclidean distance matrices between ASVs, reflecting how similarly ASVs contribute to the latent dimensions. In parallel, a functional distance matrix was derived from the binary *ASV* × *function* using Jaccard dissimilarity, which quantify the similarity between the functional profiles between ASVs. These distance matrices provide a common framework for comparison, allowing to assess to what extent the structure captured by the dimensionality reduction methods preserves functional relationships between ASVs.

To quantify this functional preservation eventually, we computed Mantel correlations between the distance/dissimilarity matrices based on functional and DRM transformed matrices. Higher Mantel correlation values indicate better preservation of functional relationships by the respective DRM-based approach, and the different approaches were subsequently ranked based on these values.

### 2.6 Topic Coherence

For both LDA- and NNMF we further computed topic coherence scores (Cv score; Röder et al. (2015)) of the derived topics. This metric quantifies the semantic consistency of the most representative features within each topic and is well-established in computational linguistics.

### 2.7 Qualitative Evaluation

Among the approaches that performed well quantitatively, we selected one to qualitatively analyze the identified topics; in contrast to the quantitative evaluation, this subsequent qualitative investigation is exploratory and aims to illustrate the types of insights that TM can reveal, rather than to provide an objective comparison across methods. The qualitative analysis focused on examining the topics’ or sub-communities’ spatial and temporal distribution and occurrence ranges for specific abiotic factors, their annotation to specific habitats and seasons as well as their taxonomic composition.

#### 2.7.1 Spatial and Temporal Specificity

To quantify the spatial and temporal specificity of topics identified by TM, we developed metrics based on the *sample* × *topic* TM output matrix. For example, the estuarine metric measures the relative prevalence of topics in estuarine environments compared to coastal or freshwater samples. To calculate this, we summarized topic contributions for estuarine (locations 2–8) and non-estuarine samples separately, normalizing by the number of samples in each category. The estuarine metric was then derived as the difference between the normalized sums of topic values for estuarine and non-estuarine samples. Formally, we calculated

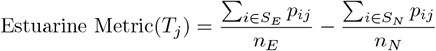

where: *T*_*j*_ denotes the specific topic, *n*_*E*_ and *n*_*N*_ represent the number of estuarine and non-estuarine samples, respectively, *p*_*ij*_ is the value of topic *T*_*j*_ in sample *i*, and *S*_*E*_ and *S*_*N*_ correspond to the sets of estuarine and non-estuarine samples.

Similar calculations were used for ‘coastal’ (sites 9-14) and ‘freshwater’ (site 1) metrics. To capture temporal patterns alongside spatial distributions, we defined four seasonal metrics: spring (weeks 12-25), summer (26-38), autumn (39-51), and winter (weeks 52 & 1-11). For comparability, values were normalized to a range of -1 to 1. This allowed each topic to be assigned a specific location and season based on the highest value, enabling the characterization of topics—or microbial sub-communities—in their occurrence and validating this pattern using the metric.

#### 2.7.2 Taxonomic Analyses

In addition to their spatial and seasonal specificity, the taxonomic composition of the topics is also part of the qualitative analysis. To enable a comparative analysis of organisms across TM-identified topics, we used our quantitatively evaluated best TM approach to create a *topic* × *ASV* matrix, which indicates the weight of each ASV in each topic. Importantly, these weights should not be equated with raw abundances in the samples as they represent how characteristic an ASV is for the co-occurrence pattern captured by a given topic. To improve comparability, this matrix was normalized across topics to importance values between 0 and 1. For each topic or sub-community, the five ASVs with the highest contributions to the respective topic were selected as a representative subset. This threshold was chosen arbitrarily to focus the analysis on the most prominent trends and differences among topics.

### 2.8 Packages

In addition to those mentioned so far, the packages Matplotlib (version 3.8.2), NumPy (version 1.26.3), Pandas (version 2.1.4), Scipy (version 1.11.4), and Seaborn (version 0.13.2) were used for data analysis and visualization (Hunter (2007); Harris et al. (2020); Virtanen et al. (2020); Waskom (2021)).

## 3 Results

### 3.1 Topic Modeling Captures Ecological Information in Low Dimensional Representations

To quantify how well the different DRM capture ecological information from the full microbiome (1551 samples × 1236 ASVs), we used Random Forest to assess their performance. We applied LDA, NNMF, PCA and PCoA transformations to generate *n*×*k* matrices (*sample*×*topic, sample*×*PCA/PCoA*−*component/coordinate*), using both unprocessed microbiome data (*none*) and preprocessed versions (*clr* and *fractions*). We evaluated these transformations across a range of *k* ∈ { 11, 21, …, 191 } and included the full-dimensional microbiome dataset as a comparative baseline (Figure S1-S9). Higher RF performance, i.e. values closer to the baseline, indicate that the respective DRM approach retained more ecological information, as captured by the respective metadata variable. For water temperature, phosphate, ammonium, nitrate, nitrite concentration, as well as the sampling week as target variables, we receive baseline performances of R^2^ or accuracy below 0.15 and thus, exclude these variables from further consideration (Table 1, Figure S1 - S6). For the remaining target variables, a noteworthy initial observation is that the highest performance was achieved with a TM in two of three cases, specifically in the prediction of chlorophyll a and the location ID (Table 1, Figure S7 and S8).

**Table 1.** Overview over the target variables, their RF performance based on the full microbiome (Preprocessed based on the best-performing preprocessing method for each case (in brackets); see Table S1) and the best DRM based RF performance. Bold target variables had a baseline performance *>* 0.15 and were further analyzed.

| Target Variable | RF performance (Full microbiome) | Best performing DRM | RF performance (Best DRM) |
| --- | --- | --- | --- |
| Temperature [°C] | $R^2 < 0.15$ | - | - |
| Phosphate [$\mu\text{mol/l}$] | $R^2 < 0.15$ | - | - |
| Ammonium [$\mu\text{mol/l}$] | $R^2 < 0.15$ | - | - |
| Nitrate [$\mu\text{mol/l}$] | $R^2 < 0.15$ | - | - |
| Nitrite [$\mu\text{mol/l}$] | $R^2 < 0.15$ | - | - |
| Calendar week | Accuracy $< 0.15$ | - | - |
| Chlorophyll a [$\text{mg/m}^3$] | $R^2 = 0.57$ (*none*) | *clr* + NNMF + $k=101$ | $R^2 = 0.62$ |
| Location ID | Accuracy $= 0.41$ (*none*) | *none* + LDA + $k=111$ | Accuracy $= 0.39$ |
| Salinity | $R^2 = 0.92$ (*fractions*) | *fractions* + PCoA + $k=41$ | $R^2 = 0.88$ |

Overall, the highest RF performance based on the full microbiome dataset was observed for salinity prediction, with R^2^ *>* 0.9. Even in low-dimensional ranges (*k* =11 and *k* =21), LDA-, NNMF-, and PCA/PCoA-based models achieved high performances (Figure S9). Predictions for location ID achieved an accuracy of 0.41 using the full microbiome dataset. TM methods (LDA with unprocessed data) approached this performance at *k* =71 and *k* =111, whereas LDA with *fractions*-transformed data performed the worst across all tested dimensions (Figure S8), Table 1).

When predicting chlorophyll a, notably, RFs trained on data derived using TM-based DRM outperformed RFs trained on the full microbiome, achieving higher performance for *k* =11 and *k* =101 (R^2^=0.58 and R^2^=0.62 respectively; Figure S7, Table 1). To explore whether other dimensionalities in the lower range near *k* =11 might also match the performance of full microbiome-based chlorophyll a predictions, we further investigated *k* values ranging between 2 and 20 in steps of 1 (*k* ∈ { 2, 3, 4, …, 20 }). Here, we focus on discussing our results with respect to the R^2^ metric, while the MAE and RMSE, which exhibit similar trends to the R^2^ metric, are provided in the SI (Table S2, Figures S10–S12). Within this range of dimensions, LDA and NNMF generally show performance comparable to PCA/PCoA, indicating that all DRM approaches maintain ecological information well when appropriately parameterized (*k*, preprocession), suggesting that multiple dimensionality reduction strategies can preserve ecological information effectively. When considering, for each approach (preprocessing + DRM), the best-performing predictions at the respective *k* value (Figure 3), remarkably, even in this low *k* range, some specific configurations (e.g., NNMF with *clr* -transformed data at *k* =8), TM-based RF models reach values slightly above the full microbiome baseline. In one TM approach, the bootstrapped mean R^2^ remained higher to the baseline, and this difference was statistically significant (Table S3). Notably, even a strong dimensionality reduction (e.g., LDA with *k* =5, a 250-fold reduction) can achieve performance close to that of the full microbiome.

**Fig. 3.**
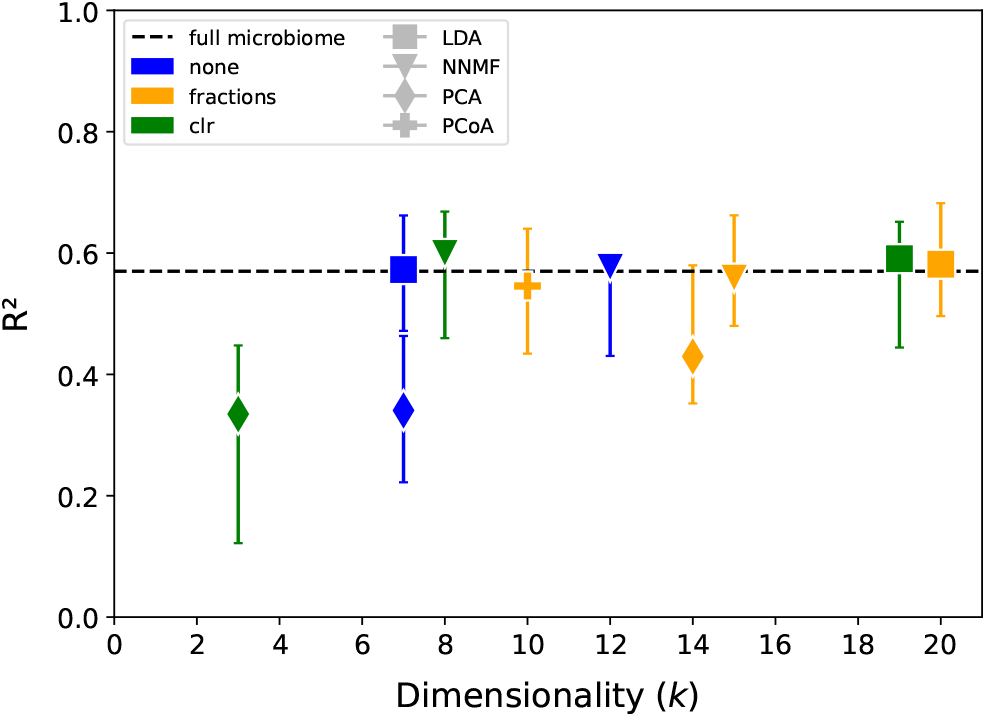
RF performance (R^2^ based on DRM generated topics or PCA/PCoA-component/coordinate across for the best performing *k* per approach across *k* ∈ { 2, 3, 4, …, 20 } when predicting chlorophyll a concentration, with 95% confidence intervals. Points represent the performance values obtained from the original RF models, while 95% confidence intervals were derived from a bootstrap procedure, like described in section 2.5.1. Only the best-performing approach (preprocession + DRM) is shown in the figure; the others can be found in Table S2 (SI).

In summary, the results of this quantitative evaluation step indicate that RF performance strongly depends on the specific target variable, with no single dimensionality reduction approach consistently outperforming the others. Additionally, the analysis revealed that even very small *k* values can be sufficient to achieve high relative RF performance. However, the prediction results for chlorophyll a—a variable representing biological structure and function—suggest that TM-based dimensionality reduction, when combined with appropriate parameter settings, allows RF models to more effectively identify patterns in environmental microbiome data compared to using the full-dimensional matrix. This was not observed with PCA/PCoA-based dimensionality reduction.

### 3.2 Topic Modeling Captures Functional Traits of the Microbiome

To further test the hypothesis that TM captures functional information in the microbiome data, we assessed how well DRM capture the ecological function of ASVs by comparing Jaccard dissimilarity of FAPROTAX-annotated functions to Euclidean distances of ASV topic or axis assignments using Mantel correlation, where higher values indicate better functional representation; to account for dimensionality effects, approaches were ranked for each *k* ∈ { 2, 3, 4, …, 20 } and averaged, where lower rank values correspond to better performance.

Our results showed that NNMF without preprocessing was the top-performing method for capturing the functional profile of the prokaryotic community (Table 2). NNMF combined with *fractions* performed nearly as well, with no significant difference from the best method. Overall, TM approaches, particularly NNMF, appear to be more effective than PCA at retaining functional information. However, we found that LDA combined with fractionated data underperformed compared to PCA-based methods, highlighting that TM can outperform PCA, but it is highly dependent on the preprocessing strategy used. These findings suggest that TM methods can be powerful tools for preserving functional relationships within microbiome data while strongly reducing dimensionality.

**Table 2.** For assessing how well the DRMs assignments of ASVs to topics or axes captures the ecological function of the ASVs, the approaches were ranked based on the average Mantel correlation between distance matrices across *k* ∈ { 2, 3, 4, …, 20 } reflecting their ability to capture the functional properties of the full-dimensional *sample* × *ASV* matrix. Lower rank values correspond to better performance. NNMF with *fractions* did not differ significantly (calculated using the Wilcoxon signed-rank test with *α*=0.05) from the best approach NNMF with no preprocessing (bold print approaches).

| Rank | Preproession | DRM | Rank value |
| --- | --- | --- | --- |
| **1** | ***none*** | **NNMF** | **1.58** |
| **2** | ***fractions*** | **NNMF** | **1.63** |
| 3 | *none* | LDA | 3.37 |
| 4 | *none* | PCA | 5.05 |
| 5 | *clr* | PCA | 5.11 |
| 6 | *fractions* | PCA | 6.16 |
| 7 | *clr* | NNMF | 6.42 |
| 8 | *clr* | LDA | 7.21 |
| 9 | *fractions* | LDA | 8.47 |

### 3.3 Topics based on NNMF and LDA with unprocessed data showed the highest coherence

To further evaluate our models, we calculated topic coherence scores (Cv) for our TM methods LDA and NNMF across the lower *k* range (*k* ∈ { 2, 3, 4, …, 20 }). Topic coherence quantifies how consistently the top ASVs within a topic co-occur, with higher values indicating more internally consistent, interpretable topics. We found that LDA and NNMF applied to unprocessed microbiome data consistently achieved the highest Cv values across the examined *k* range. Other approaches showed lower and more variable coherence scores, with NNMF applied to *fractions*-processed data performing relatively best among them (Figure S12).

The results presented to this point show that TM is effective in capturing ecological and functional relationships in a microbiome from a highly dynamic environment, provided the right combination of preprocessing and TM algorithm is applied. Furthermore, our preliminary analyses indicate that there is not one single, universally optimal *k* value. This finding underscores the need for multi-scale approaches, when applying TM to microbiomes, to effectively capture various levels of community structure.

### 3.4 TM-identified Topics Show Clear Spatial and Temporal Patterns

DRM are commonly used in exploratory data analysis to uncover relevant latent patterns in large datasets. Thus, aside from the quantitative measure of how well a DRM maintains relevant information, the interpretability of its output is central for its usefulness. Because interpretability is inherently qualitative, it is not suitable for a comparative study across methods. Therefore, for our qualitative investigation, we focused on a single DRM that performed well in our quantitative evaluation—a NNMF model with *k*=11 applied to unprocessed microbiome data. This choice was motivated by the fact that, during our quantitative evaluation, *k*=11 represented one of the lowest values at which the RF performance of the TM (using NNMF with unprocessed data) first exceeded that of the full-dimensional *sample* × *ASV* matrix. Furthermore, NNMF applied to the unprocessed microbiome data captured the highest level of functional information from the microbial community. Additionally, for NNMF on the unprocessed data, the topic coherence values remained high across all evaluated k-values, which further supported our choice.

With this chosen approach, we visualized the topic distribution across samples, revealing distinct spatial and temporal patterns (Figure 4A). For example, Topic 2 is mainly occurring at location 1, the freshwater location, for the whole year. In contrast, Topics 1, 3, and 6 are primarily concentrated in coastal locations, each exhibiting different seasonal occurrence patterns. In the estuary, Topic 4, 5, 7, 8 and 11 are predominant - also with different temporal occurrence foci. Topic 4 is mainly occurring in summer and autumn, while Topic 5 has a very narrow time range and appears more frequently in summer. The latter also applies to Topic 7, while Topic 8 and Topic 11 are most frequently observed within a very narrow time frame during spring resp. autumn. To further formalize the spatial distribution, we calculated the specificity of the topics with regard to the three habitats covered by our dataset (Figure 4B). This metric-based approach supported the previous observation that the Warnow Estuary hosts five sub-communities (assemblages of co-occurring ASVs) with the highest specificity for this dynamic habitat, each exhibiting distinct temporal patterns (Topic 4: summer, autumn; Topic 5: summer; Topic 7: summer; Topic 8: spring; and Topic 11: autumn). Notably, no distinct estuarine sub-community was dominant in the estuary during winter or persisted year-round, as was observed for the coastal and freshwater sub-communities (Figure 4A).

**Fig. 4.**
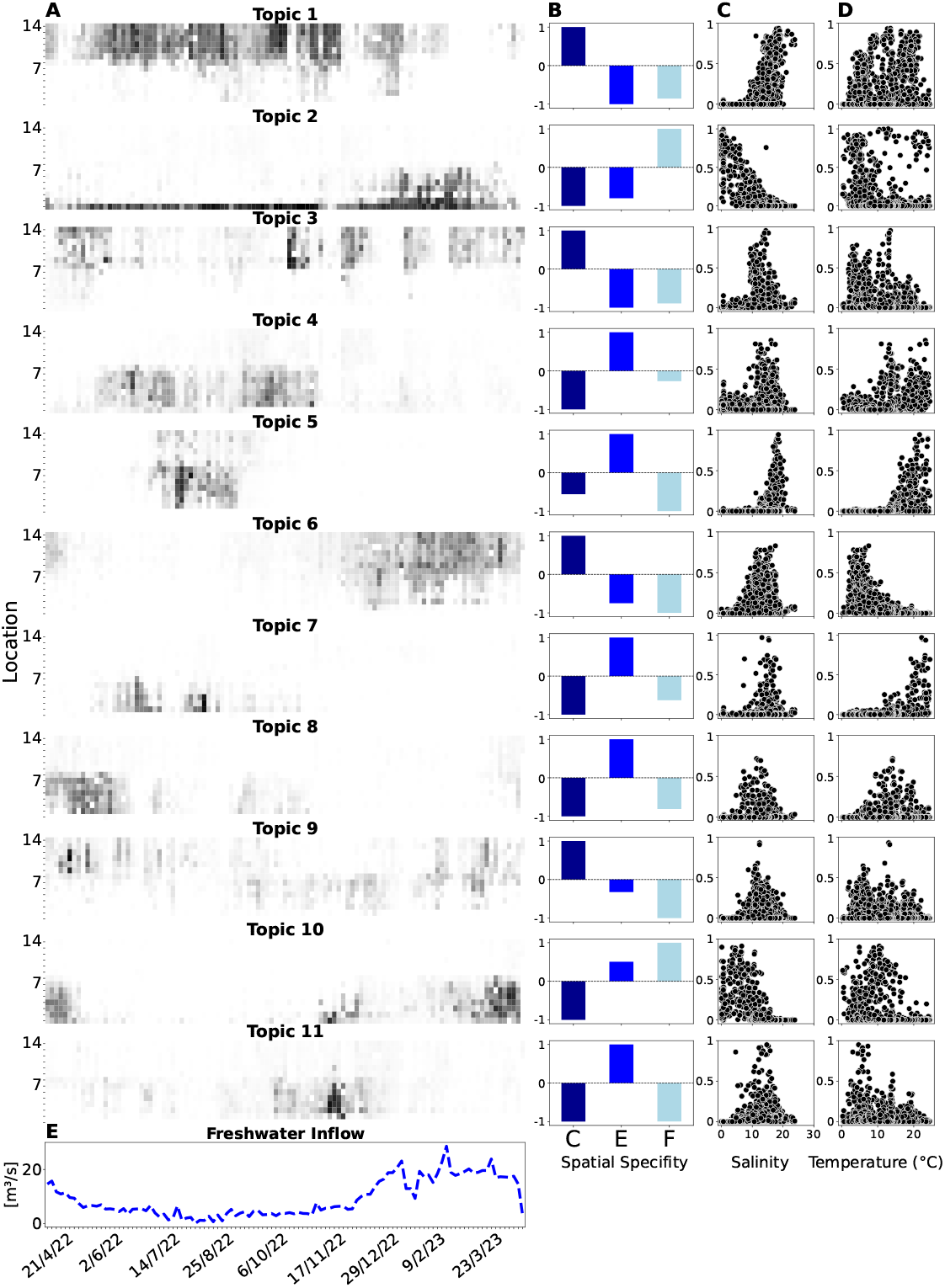
**A**: Applying the evaluated approach of NNMF with a *k* =11 on unprocessed microbiome data led to the identification of 11 microbial sub-communities (assemblages of co-occurring ASVs) showing a clear spatial and temporal pattern. Heatmaps display topic abundances per sample, ranging from 0 to 1, with darker shading indicating higher abundances. Sampling locations comprise one freshwater site (Location 1), estuarine sites (Locations 2–8), and coastal sites (Locations 9–14) (compare figure 2). **B**: The spatial specificity of these sub-communities—categorized into coastal (C, dark blue), estuarine (E, blue), and freshwater (F, light blue) environments—was assessed using spatial and seasonal metrics (compare section 2.7.1). The higher the metric value, the higher the specificity for a spatial category. This analysis identified distinct estuarine sub-communities (Topics 4, 5, 7, 8, and 11). Each sub-community exhibits a unique observed association with specific salinity and temperature ranges, as illustrated by the topic loading per salinity/temperature (**C, D**). The y-axis in the salinity and temperature scatterplots represents the topic loadings per sample. **E**: The blue line indicates the freshwater inflow volume into the estuary [m^3^/s] (Inflow data originate from one sampling site downstream of the estuary on the Warnow River), appearing to influence the distribution of these sub-communities.

These findings demonstrate that TM effectively identifies sub-communities highly specific to certain habitats and times of the year.

### 3.5 Seasonal Succession of the Estuarine Topics

Notably, the temporal pattern of the TM-identified distinct estuarine sub-communities resembles a seasonal succession: in spring, Topic 8 appears at all estuarine sites in high abundances, followed by the sub-communities Topic 4 and Topic 7. Topic 7’s presence is briefly interrupted as it is replaced by sub-community Topic 5, which initially appears at the estuarine stations closer to the Baltic Sea, while Topic 7 reappears in the southern estuarine zones. Both sub-communities gradually diminish with the approach of autumn, eventually giving way to Topic 11, which peaks in late November before disappearing quickly again. Prior to Topic 11’s peak, sub-community Topic 4 also fades out. At this point in time, the distribution heatmap shows the presence of freshwater sub-communities Topic 2 and Topic 10, as well as the coastal sub-community Topic 6 for the estuarine areas (Figure 4A).

To further investigate this succession, we incorporated various environmental variables. Specifically, we linked the topic loadings to salinity and temperature, both known to influence the structure and distribution of microbial communities (Herlemann et al. (2011), Lozupone and Knight (2007), Liu et al. (2012), Kirchman (2008), Rivkin et al. (1996)) (Figure 4C, D), as well as to the freshwater inflow volume into the estuary (Benson (1981); Alber (2002)). In the Warnow Estuary, this inflow primarily originates from the Warnow River (Figure 4A, blue line).

With a relatively broad distribution range for salinity (8–15) and temperature (8–17°C), Topic 8 appears in the estuary during spring, when freshwater inflow is declining and at a low level. Later, Topic 4, which shows an observed range of 10–20°C, enters the estuary and remains dominant until autumn. Its persistence is likely due to its tolerance also for lower temperatures, as it appears as well at temperatures below 5°C. Shortly after Topic 4, Topic 5 and Topic 7 follow—both sub-communities are associated with high-temperature (*>*20°C) and a limited occurrence in summer. During this period, freshwater inflow into the estuary remains low. Since both Topic 5 and Topic 7 have a relatively narrow and high saline distribution range (15–20), this low freshwater influence, encoding declines in salinity, appears to be conditional for their dominance in the estuary at this time.

As autumn approaches, freshwater inflow begins to increase. Around this time, Topic 4 disappears, and Topic 11 becomes dominant in the estuary. Despite its association with low temperatures (2-5°C), Topic 11 quickly vanishes, while freshwater inflow continues to rise. The resulting decline in salinity suggests why two freshwater-associated sub-communities, Topic 2 and Topic 10, become dominant in the estuary after Topic 11 with its higher salinity requirements (around 10) disappears. Meanwhile, in the upper estuary, the coastal sub-community Topic 6 becomes more prominent.

During this transition, the distributional patterns of Topics 2, 6, and 10 resemble a well-coordinated puzzle. Initially, Topic 2 dominates the lower estuary, while Topic 6 prevails in the upper estuary. However, as spring arrives, the cold-adapted Topic 6 (Figure 4A, C, D) is displaced by Topic 10, which emerges just as Topic 2 disappears from the estuary.

Overall, our findings indicate that the succession of sub-communities in the estuary is associated with their occurrence across specific temperature ranges. However, we observed that under prolonged and pronounced salinity changes, driven by increased freshwater inflow, the established estuarine sub-communities could not persist and were gradually displaced by freshwater and coastal communities.

### 3.6 Distinct Taxonomic Composition of Estuarine Sub-communities During Their Seasonal Succession

The previously gained discovery of distinct estuarine microbial sub-communities and their seasonal succession was derived from the analysis of the *sample* × *topic* matrix generated by our evaluated TM approach. To gain insight into the changing structure of taxonomic sub-communities, the TM approaches used here also produce a *topic* × *ASV* matrix, which provides the contribution of each ASV to each topic. It should be noted that, the weights in this matrix do not represent absolute or relative abundances of ASVs in the samples, but rather indicate how characteristic a given ASV is for a particular topic. To explore how these taxonomic patterns of TM-identified estuarine sub-communities change during their seasonal succession, we considered the five ASVs with the highest contributions to each topic and examined their contributions across the other topics as well (Figure 5).

**Fig. 5.**
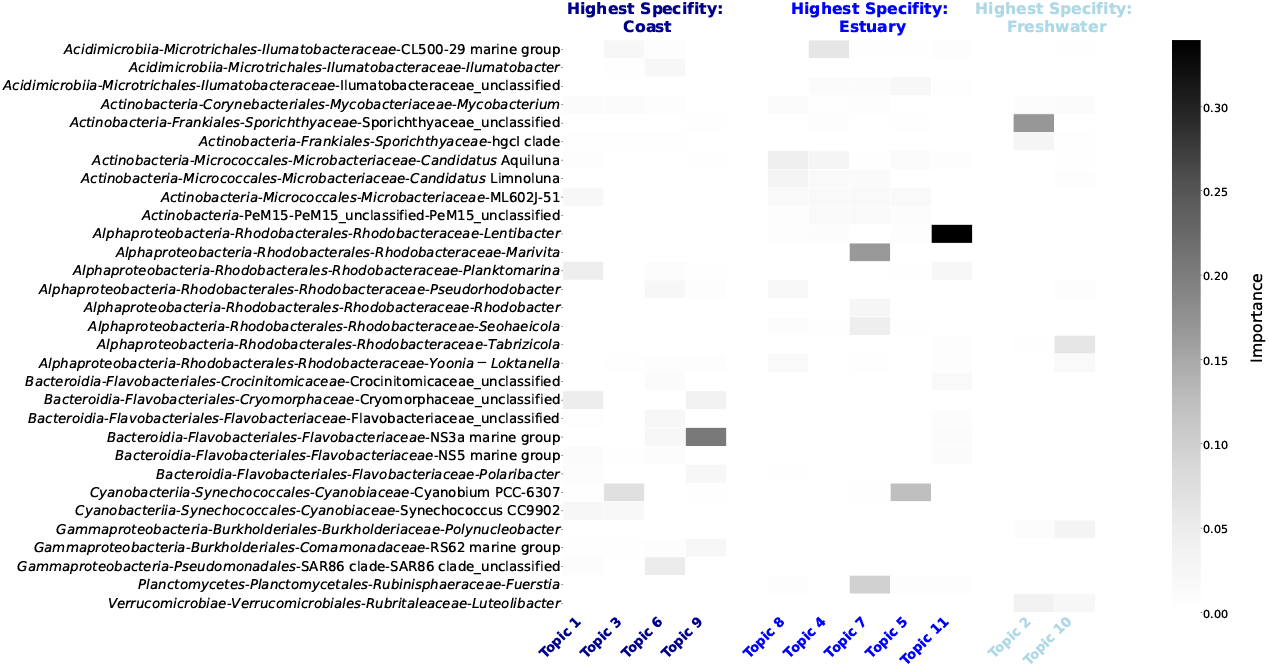
Distribution of the top 5 contributing ASVs per topic (or microbial sub-community) and the contribution of these organisms across the others (coastal: dark blue, estuary: blue, freshwater: light blue). The taxa are listed in the format: bacterial class - order - family - genus (IDs in Table S4). The intensity of the heatmap fields indicates the contribution value the ASV had toward the definition of the topic, based on the TM output *topic × ASV* matrix.

These data indicate that different *Actinobacteria* contribute substantially to almost all estuarine sub-communities (Topics 8, 4, 7, and 5), while they were barely represented among the most important ASVs in coastal and freshwater topics. However, in estuarine Topic 11 only a single *Actionbacteria, Candidatus* Aquiluna, was slightly contributing. Instead, representatives of *Bacteroidia*—a class mainly important here within coastal associated sub-communities—were more important, while being not the main contributors in the other estuarine topics.

Regarding the seasonal succession of the estuarine sub-communities, we observe a clear shift in the dominant organisms: In spring, the sub-community described by Topic 8 is specifically characterized by *Rhodobacteraceae*-belonging genus *Pseudorhodobacter*, which belong to *Alphaproteobacteria*. Topic 4 follows with a predominance of *Acidimicrobiia*, specifically the genus CL500-29 marine group belonging to the family *Ilumatobacteraceae*. Topic 7 is dominated by two important organisms, one belonging to the *Rhodobacteraceae* (*Alphaproteobacteria*) family, namely *Marivita* and the other one to *Rubinisphaeraceae* (*Planktomycete*s), called *Fuerstia*. Sub-community Topic 5 is dominated by Cyanobium PCC-6307 from the family *Cyanobiaceae*, a family within the *Cyanobacteriia*. In sub-community Topic 11, *Rhodobacteraceae* (*Alphaproteobacteria*) is again the most contributing group, but here the genus *Lentibacter* predominates.

### 3.7 How Topic Dimensionality Shapes the Resolution of Estuarine Sub-communities

The results presented in the previous exploratory section were produced at *k* =11. In order to show that the finding of a succession in the estuary is robust against the choice of *k*, we analyzed the seasonal and spatial distribution of topics across values of *k* from 3 to 12 (Figure 6).

**Fig. 6.**
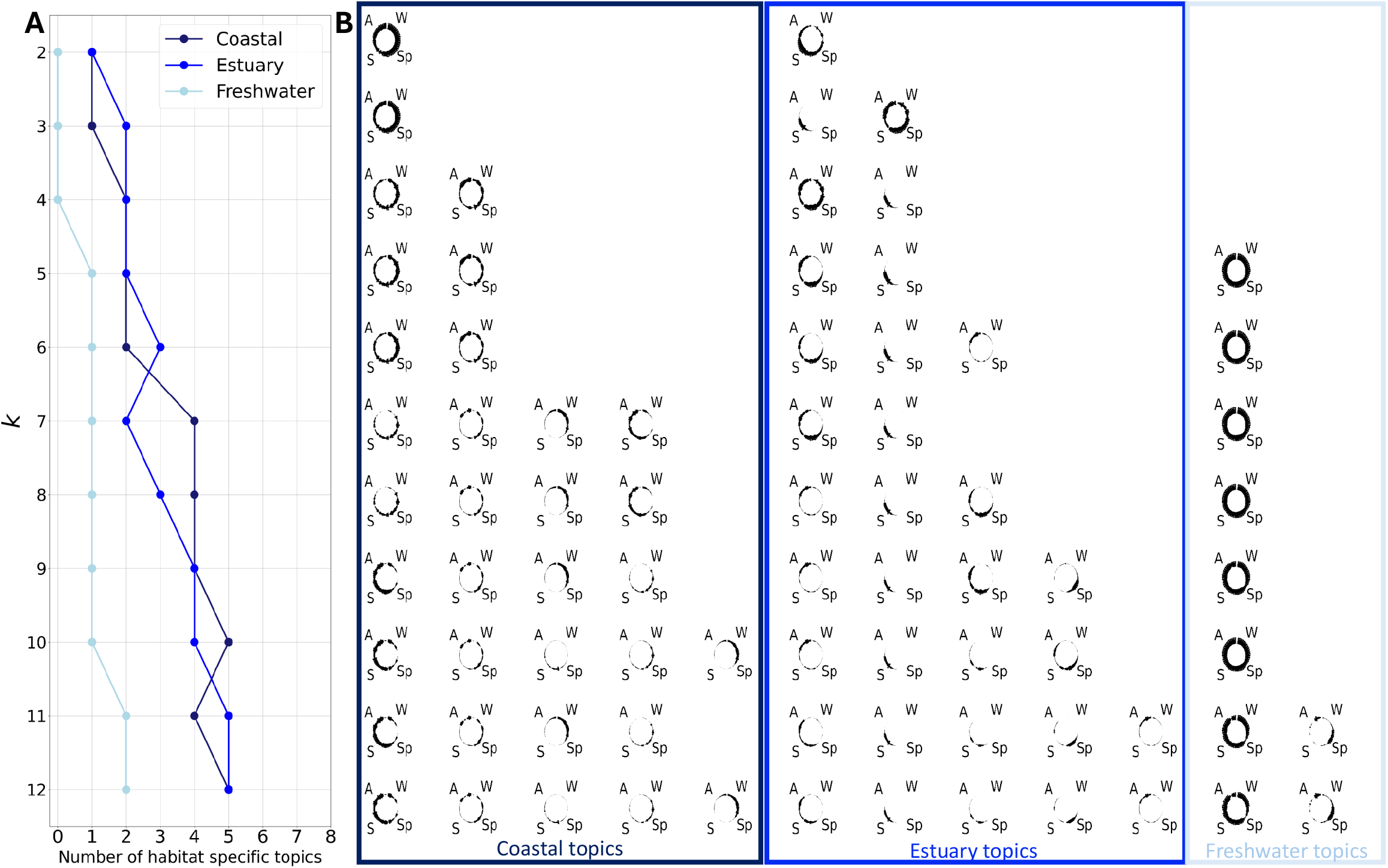
Differences in spatial and seasonal distribution of topics over a range of *k* (*k* ∈ { 2, 3, 4 }). **A**: The number of habitat-specific topics (coastal: dark blue, estuary: blue, freshwater: light blue) at different *k* values with their **B**: seasonal distribution (Spring (Sp), Summer (S), Autumn (A), Winter (W)). From *k* =6, the topics can be classified into the 3 different habitat types for the first time. Over the *k* range, there are always fewer topics that are specific to freshwater. Up to *k* =10, there is only one freshwater topic that occurs almost all year round. In contrast, the higher the *k* is set, the more temporal nuances of the coastal and estuary-specific topics become visible.

Examining the distributions for *k* ∈ { 2, 3, 4 }, it is evident that no freshwater-specific topic can be identified. However, within the estuary, these dimensionalities reveal a year-round estuarine distinct sub-community, which is no longer present at higher *k* values. The first clear distinction between estuarine and freshwater sub-communities appears at *k* =5, marking a threshold where the dimensionality is sufficient to differentiate these two habitats. This suggests that *k* =5 represents the minimum complexity required for our data to meaningfully extract estuarine sub-community dynamics distinct from those of freshwater environments.

Between *k* =2 and *k* =7, the number of topics associated with the estuary fluctuates between 2 and 3. This number increases to 4 at *k* =9 and ultimately reaches 5 at *k* =11. The progressive addition of topics aligns with a growing ability to resolve more detailed seasonal differences. For example, at *k* =6, we observe the emergence of topics associated with spring and early summer, summer, and fall, revealing the dynamic shifts in microbial community structure over these periods. By *k* =9, a distinct spring topic is distinguishable, indicating the model’s capacity to capture even finer temporal nuances at higher dimensionality.

These results emphasize that different *k* values provide distinct perspectives on the data. While lower *k* values reveal dominant patterns and overarching trends, higher *k* values offering a more nuanced distinction of habitat- and time-specific sub-communities, with temporal succession already becoming apparent at *k <* 11.

## 4 Discussion

A comprehensive understanding of microbial communities can facilitate more accurate detection of ecosystem changes and, consequently, the development of appropriate conservation measures. Gaining such an understanding is particularly challenging in highly dynamic ecosystems due to multivariate and often unpredictable changes. This complicates the monitoring of potentially harmful alterations, such as those caused by anthropogenic influences (Elliott and Quintino (2007)). In this study, we investigated the utility of the unsupervised ML technique topic modeling—a method primarily used in natural language processing—to elucidate microbial dynamics in a highly dynamic system, using the Warnow Estuary and its surrounding area as a case study.

### 4.1 Enhanced Ecological and Functional Insights Through TM-Based Data Transformation

In linguistics, the original application domain of TM, evaluating TM-generated word clusters is relatively straightforward, as semantic relationships between grouped terms can often be easily interpreted. In contrast, the evaluation of the meaningfulness of microbial topics is more complicated. While some TM methods, such as LDA, provide internal evaluation metrics (e.g., LDA perplexity), we adopted a model-independent evaluation approach that also incorporates ecological variables. This allows for a simultaneous comparison of different DRM applied to various pre-processed datasets across a broad range of *k* values while maintaining ecological context. To achieve this, we employed RF as a supervised, external evaluation tool. By comparing RF prediction performance based on the full microbiome dataset with that of the DRM-derived matrices, we assessed the extent to which ecological information could be captured using TM- and PCA/PCoA-based dimensionality reduction approaches.

This analysis yielded several key insights. First, certain target variables could not be effectively predicted by RF models. Notably, water temperature—as a described influence factor on microbial communities, a commonly measured abiotic variable in ecological studies (Kirchman (2008), Rivkin et al. (1996))—consistently exhibited poor predictive performances across all dimensionality levels. We attribute this underperformance primarily to the temporal train-test split in our RF model setup. Specifically, the model was trained on data from April to January and tested on data from January to April. Given that water temperature exhibits strong seasonal fluctuations, this temporal split likely impaired the model’s predictive accuracy. For salinity, chlorophyll a, and the location ID, RF achieved solid predictive performance, suggesting that the bacterial community composition has a strong correlation with these factors.

A second key finding during the quantitative evaluation, was that the highest RF R^2^ value was achieved for salinity and exceeded 0.9 when using the full microbiome. While the predictive performance of TM- and PCA/PCoA-based RF performances was consistently lower than that performance based on the full dimensional dataset, it remained at a very high level overall. This strong performance confirms a robust relationship between bacterial community composition and salinity in our highly variable study habitat—a relationship that has been described for other habitats before (Herlemann et al. (2011), Lozupone and Knight (2007), Liu et al. (2012)). The RF based on the TM representations did not outperform the full dataset based performance, but they offered complementary ecological insight: topics grouped ASVs into sub-communities restricted to narrow salinity ranges (Figure 4C).

Performing the RF evaluation in general, we expected that the reduction of dimensions, i.e., the intentional removal of data features, would result in a loss of information and that RF models based on DRM-transformed data would perform worse than those based on the full dimensional dataset (Sperlea et al. (2022)). The next key finding, in predicting chlorophyll a, was the striking performance of multiple RF models based on TM methods (LDA and NNMF), which achieved higher predictive accuracy using remarkably fewer dimensions than models trained on the full microbiome. We identified multiple approaches where RF performance either exceeded or closely matched that of the full-dimensional microbiome, depending on the choice of *k* - suggesting that RF, when using TM-clustered units, captures a comparably detailed or even more refined representation of the full community when predicting chlorophyll a levels. The fact that this occurs across multiple *k* values confirms that important patterns in community distribution do not emerge at a single “optimal” *k*, but rather across different *k* values, underscoring the inherently multi-scale nature of microbial community structure. To further support this, figure 6 shows that from *k* =5 onward, freshwater and estuarine communities were clearly separated, representing the minimum level of complexity needed to meaningfully capture estuarine community dynamics in comparison to freshwater environments. This is underpinned by the fact, that at *k* =5, near-optimal performance was also achieved in the TM-based RF prediction of chlorophyll a. These findings highlight that different *k* values provide distinct and complementary perspectives on microbial community structure. Lower *k* values capture broad patterns and dominant trends, such as major successional shifts, whereas higher *k* values reveal finer seasonal and spatial variations. Although the choice of the parameter *k* can and should be supported by data, the final choice of *k* ultimately remains somewhat arbitrary, as there is currently no method that determines the optimal *k* for every problem. It is therefore crucial that this decision is tailored to the research question, bearing in mind that different values of *k* may provide complementary ecological insights rather than a single “true” solution, and that it may be necessary to analyze topics with different *k* values.

In addition, when proceeding to the second quantitative evaluation step, we discovered that NNMF performed best when applied to unprocessed or fractionated data for capturing the functional properties of the microbiome. Regarding the functional evaluation, a few limitations must be noted. For one, the FAPROTAX-derived functional potential used in this study is based on the genomic presence of genes and, thus, does not necessarily reflect the actual functional activity that would be reflected in the expression of these genes. Furthermore, the accuracy of FAPROTAX-based predictions depends heavily on the availability and quality of ASV-specific information. In our microbiome dataset, this information was available for only about one-third of the ASVs. Consequently, our predicted functional distributions should be interpreted as exploratory approximations rather than exact representations of ecological function.

Notably, during the course of the entire quantitative evaluation process, it became apparent that TM often performed well on unprocessed microbiome data. This result is counterintuitive because of the compositional nature of ASV data and because the sample-specific sequencing depth is, to a large degree, a technical artifact. Nevertheless, this finding may be explained by the fact, that both LDA and NNMF are designed to operate on non-negative input and yield additive, interpretable components. LDA explicitly represents samples as probability distributions over latent topics, while NNMF implicitly identifies latent factors with probability-like loadings. In this way, both methods match the generative logic of compositional systems, where each sample is naturally a mixture of overlapping latent sub-communities. In contrast, linear approaches such as PCA allow negative loadings and enforce orthogonality of components. Thus, the performance of LDA and NNMF on raw microbiome data may reflect their alignment with the underlying data structure highlighting their practical value as complementary tools for exploring microbial community structure.

Overall, while TM demonstrated advantages in capturing complex microbial community patterns and reducing the dimensionality of microbiome data, inherent limitations of TM should be acknowledged. In the probabilistic model LDA, the TM-inherent bag-of-words assumption (Blei et al. (2001)) treats each sample as a collection of independent features, which may obscure biologically meaningful relationships between taxa. In addition, TM methods such as LDA and NNMF naturally lack explicit phylogenetic awareness. While taxa with similar ecological distributions are often grouped together due to their common occurrence in the samples, subtle differences in distribution patterns can still lead to their division into multiple topics, especially when a higher number of topics is used, resulting in a finer division into subcommunities. This should be kept in mind, even though TM performed robustly in our quantitative evaluation when appropriately parameterized (in terms of preprocessing and the number of topics).

Moreover, it is important to consider the conceptual interpretation of topics: TM derived topics are not equivalent to well-defined ecological entities, like communities. However, while topics cannot be directly linked to established ecological concepts, such as guilds, they share certain similarities. Like guilds, topics capture groups of co-occurring taxa that may reflect functional or ecological roles. Our analysis of predicted functional distributions suggests that functional information is, at least partly, embedded in the topic structure, although not in a one-to-one correspondence with classical ecological definitions. However, a key difference is that the composition and number of topics depend on the chosen parameter *k*, whereas ecological concepts like guilds (Simberloff and Dayan (1991)) are theoretically independent of such modeling choices. Thus, we view topics as data-driven abstractions resembling guild-like structures with great use in exploratory data analyses; further validation with longer-term or experimental data would be needed in order to show whether these abstractions are fully complementary with theoretical concepts.

### 4.2 TM as an Additional Tool for PCA/PCoA-based Approaches in Microbiome Studies

We found that the highest RF performance based on DRM was achieved using a TM approach rather than PCA/PCoA for the prediction of chlorophyll a and the location ID (Table 1). PCA identifies axes of greatest variance, requiring them to be both linear and uncorrelated. While effective for reducing dimensionality, this constraint may prevent PCA from detecting subtle ecological relationships, particularly when interactions between the variables are nonlinear. In contrast, TM identifies latent ecological structures by grouping features based on shared patterns of occurrence rather than purely on variance. This provides a complementary perspective, offering an alternative way of revealing potentially ecologically meaningful clusters that may not be captured by variance-based methods. However, some RF models based on PCA performed well, particularly in predicting salinity (Figure S9). Moreover, PCA has demonstrated robust performance in other biological studies, including research on the gut microbiome (Wakita et al. (2018), Liu et al. (2020)). For salinity prediction, the RF model achieved its highest performance when using PCoA transformed data units. This supports the suitability of PCoA—commonly applied in microbiome research—for identifying ecologically meaningful low-dimensional clusters (Paliy and Shankar (2016)). However, in contrast to TM, PCoA has the significant limitation of not providing feature distributions—meaning it does not indicate which ASVs are relevant within the generated subgroups. However, both PCA and PCoA have well-established strengths—such as computational efficiency and the generation of interpretable linear axes making them highly valuable tools for microbiome research. With our results, we propose incorporating TM as a complementary dimension reduction method that can provide additional insights alongside PCA/PCoA. Promising new approaches for dimension reduction of microbiome data while capturing nonlinear relationships come in the form of deep representation learning approaches (Oh and Zhang (2020)). However, as of now, these are not available for environmental microbiome data.

### 4.3 Identification of Estuarine Communities and Their Seasonal Dynamics

A central goal of ecology is understanding what determines the distribution and abundance of communities (Cody and Diamond (1975), MacArthur (1984)). Using TM with the settings quantitatively evaluated in the first part of this work (NNMF, unprocessed data, *k* =11), we explored the microbiome dataset qualitatively to identify characteristic spatial and temporal patterns.

We identified five distinct estuarine sub-communities displaying an annual succession: Topic 8 emerged in spring, followed by the appearance of Topics 4 and 7. Topic 7 was briefly replaced by Topic 5, which gradually diminished as autumn approached. By late November, Topic 11 peaked before disappearing rapidly. Just before Topic 11 reached its maximum abundance, Topic 4 also began to fade. In the first half of December, Topic 11 vanished entirely, and freshwater and coastal sub-communities alternated in dominance across different parts of the estuary.

We explain this succession through the observed specific salinity and temperature ranges that enable these sub-communities to exist and dominate at particular locations and times when conditions are optimal for them (Figure 4). Previous studies suggest that communities in transitional zones emerge as a fusion of freshwater and marine microorganisms, with some taxa disappearing while others, specialized for transitional conditions, thriving (Rocca et al. (2019)). When specific habitat parameters, like salinity, shift permanently, more competitive communities once again dominate the transitional area. This is suggested by our analysis when considering Topic 2, Topic 10, and Topic 6, particularly when freshwater input is on a constant high level. Their concurrent dominance in freshwater and coastal areas during this period indicates that they did not migrate into the estuary due to displacement but instead expanded opportunistically as environmental conditions became favorable.

In addition to the observed seasonal succession of estuarine sub-communities, an important observation was the absence of an estuarine sub-community that persisted year-round or exclusively occurred during winter—a phenomenon observed in the coastal and freshwater sub-communities (Figure 4A). While estuarine sub-community Topic 11 did exhibit broad temperature tolerances, potentially allowing to persist in winter, this finding suggests that temperature alone cannot explain the observed seasonal patterns. Considering the observed specific salinity ranges in which the the sub-communities’ exists alongside an additional parameter—the volume of freshwater inflow, which correlates with long-term salinity changes—added an additional layer of interpretability. The seasonal patterns were primarily driven by fluctuations in freshwater inflow, reinforcing the strong relationship between salinity and bacterial community structure revealed by our RF-based analysis (Figure S9).

It is important to note, however, that these patterns were inferred from a single year of sampling. Consequently, the observed succession and apparent environmental associations may reflect short-term dynamics specific to this year rather than consistent ecological processes. Longer-term time series would be necessary to verify whether these patterns are robust and representative of recurring estuarine dynamics, or whether they instead reflect transient or year-specific conditions.

#### 4.3.1 Taxonomic Composition of Estuarine Sub-communities Changes During Their Succession

For the observed seasonal succession in the TM identified estuarine sub-communities, we describe the key taxa associated with each sub-community using TM (Figure 5), noting that only the top five ASVs per topic are shown to illustrate broad trends. Furthermore, it should be noted, that topics reflect co-occurrence patterns rather than relative abundances, prominent ASVs of a certain topic, such as *Actinobacteria* in the estuarine topics, are not necessarily exclusive to these sub-communities. Therefore, for supporting the biological interpretation of the TM derived topics, follow-up studies could include additional analyses that link highly weighted ASVs to their distribution across samples (e.g., patterns of presence and absence or differences in abundance between samples with high and low topic abundances).

In spring, sub-community Topic 8 with its important family *Rhodobacteraceae* (*Alphaproteobacteria*), including the genus *Pseudorhodobacter* appears. This genus has previously been described as a key component of Baltic Sea microbial communities during spring (Bunse et al. (2016)), aligning with our temporal observations. It has also been recognized as an important taxon in previous Baltic Sea studies (Herlemann et al. (2011)).

Following Topic 8, Topic 4 became dominant in the estuary, characterized by the CL500-29 marine group of the family *Ilumatobacteraceae* (*Acidimicrobiia*). This taxon has been identified in winter microbial communities in the Pearl River Estuary (Zhang et al. (2023)). Here, the broad seasonal presence of Topic 4 in the Warnow Estuary suggests that its members might be temperature-tolerant organisms, potentially explaining their persistence across multiple seasons.

Topic 7, emerging earlier in the summer than Topic 5, is defined by the alphaproteobacterium *Marivita* and the planctomycete *Fuerstia*. These taxa play distinct ecological roles: *Fuerstia* has been described as a macrophyte associate in the Baltic Sea and was also found in the German Wadden Sea, another dynamic habitat. *Marivita* has been linked to cyanobacterial blooms, particularly of *Nodularia spumigena* (Munkes et al. (2021), van der Loos et al. (2023), Kohn et al. (2016)).

As the summer progresses, Topic 7 is displaced by sub-community Topic 5, which represents a summer-dominant estuarine sub-community constrained to narrow salinity and temperature ranges. The primary taxon in Topic 5, the cyanobacterium Cyanobium PCC-6307, is well-adapted to high-light summer conditions and has been frequently reported in Swedish freshwater blooms, the Baltic Sea, and brackish estuarine regions (Dirks et al. (2024), Zhang et al. (2023), Larsson et al. (2014)). Studies suggest that Cyanobium PCC-6307 combines traits of marine and freshwater lineages, representing a brackish water-adapted genotype that is particularly prevalent in summer (Larsson et al. (2014)). This aligns with previous findings describing the coexistence of freshwater and saltwater bacterial strains in brackish systems such as the Baltic Sea (Herlemann et al. (2011)).

Cyanobacterial blooms in the Baltic Sea are common in summer (Munkes et al. (2021)). The alternating dominance of Topic 5 and Topic 7 may indicate that favorable conditions in early August promote the prevalence of Cyanobium PCC-6307 (Topic 5). In contrast, late July and late August might favor the dominance of *Nodularia spumigena*, which is indirectly indicated by the prevalence of *Marivita* in Topic 7.

Finally, as Topic 4 gradually fades out, Topic 11 begins to increase in prevalence within the estuary. This sub-community is dominated by *Lentibacter*, an alphaproteobacterium in the family *Rhodobacteraceae*, which thrives under estuarine-specific conditions before the influx of freshwater. Previous studies have associated *Lentibacter* and the NS5 marine group with estuarine systems, such as the Dagu River estuary, where *Lentibacter* showed higher abundances during winter (Yang et al. (2022)), aligning with our results.

#### 4.3.2 *Actinobacteria* in Estuarine Sub-communities

Figure 5 shows that the composition of the distinct estuarine sub-communities differs markedly from those associated with coastal and freshwater environments. Notably, *Actinobacteria* representatives are important in the estuarine topics, with the exception of Topic 11, where this group is largely absent apart from a low contribution of *Candidatus* Aquiluna. The *Actinobacteria* genera identified as important in the other estuarine sub-communities include besides *Candidatus* Aquiluna, *Candidatus* Limnoluna, and ML602J-51. These taxa have also been reported in estuaries such as the Bailang River Estuary, Arctic fjords, and the Parker River Estuary (Dong et al. (2023), Kang et al. (2012), Crump et al. (2004)).

The genus *Candidatus* Aquiluna, present in all estuarine sub-communities, has been positively correlated with ammonium and dissolved inorganic nitrogen concentrations, as well as with eutrophication levels in the Beibu Gulf. It has also been linked to eutrophic water conditions in other studies (Li et al. (2020b), Hahn (2009)). Given that the Warnow River and its estuary are considered highly eutrophic, this likely provides a favorable environment for this organism (Freese et al. (2006), Warkentin et al. (2011), Rönspieß et al. (2020)).

In the northern Baltic Sea, *Actinobacteria* abundance has been described as temperature-sensitive, increasing with higher temperatures (Holmfeldt et al. (2009)). This aligns with our observation that Topic 11, which occurs during the cooler temperatures of late autumn, contains only one low important representative of *Actinobacteria*. During this time, the niches of *Actinobacteria* may be occupied by other organisms, such as *Bacteroidia*, which were important in the estuarine sub-community described by Topic 11. Within *Bacteroidia*, a class absent in other estuarine communities but prevalent in coastal topics, genera such as the NS3a marine group and NS5 marine group are important. These have been described as common in the microbial communities of the Baltic Sea before (Lindh et al. (2016)).

Overall, evidence from other environmental studies suggests that the seasonal occurrence of key taxa observed here is plausible, supporting the idea that the classification of organisms by TM, in this case NNMF, may provide useful insights. In this way, TM could serve as a complementary approach alongside taxonomic studies.

## 5 Conclusion

In this study, we investigated whether TM can help make a high-dimensional microbiome dataset more interpretable on a community level, thereby improving our understanding of a highly dynamic habitat, using the Warnow Estuary as a complex test case. To achieve this, we applied a two-step quantitative assessment process using first a supervised ML method, incorporating ecological variables to compare TM and PCA/PCoA in different set ups in terms of their ability to preserve ecological information. In the second step, we examined the preservation of functional information after applying TM and PCA. These analyses indicate that TM performed comparably to, and in some aspects may better capture, microbiome information than conventional PCA in our dataset. While PCoA also showed strong performance for several target variables, it lacks a key advantage of TM: the ability to provide interpretable feature distributions. In addition, we evaluated a TM approach that proved to be well suited for our microbiome data in our quantitative evaluation and conducted a qualitative assessment. This evaluation provided important exploratory insights into the microbial community of the Warnow Estuary. We identified five sub-communities that exhibited the highest specificity for the estuarine environment and displayed a clear seasonal succession for the sampling year, partially explained by freshwater inflow from the Warnow River. Each of these sub-communities had a distinct salinity and temperature distribution spectrum and exhibited unique compositions of key taxa, with *Actinobacteria* occuring in all estuarine sub-communities.

In summary, our work demonstrates that TM methods, originally developed for a completely different application domain, can be valuable supplementary tools for analyzing environmental microbiome datasets and enhancing our understanding of highly dynamic habitats such as the Warnow Estuary. Building on this, TM could serve as a foundation for ecosystem state assessments, including those of highly dynamic and anthropogenically influenced environments.

## Supplementary information

The Supplementary Information can be found in TM on microbiome SI.pdf.

## Declarations

### Ethics approval and consent to participate

Not applicable.

### Consent for publication

Not applicable.

### Availability of data and materials

Demulitplexed and primer-clipped read data are available from the European Nucleotide Archive (ENA) with MIxS compatible metadata using the data brokerage service GFBio and are available under the study accession numbers PRJEB88008 (16S) and PRJEB88011 (18S). Contextual data is publicly stored in the IOWDB and can be accessed via the DOI 10.12754/data-2025-0001. All utilized functions and their application for the generation and evaluation of topics or PCA and PCoA component/coordinate are available in the GitHub repository: https://github.com/a-k-29/topic_modeling_microbiome.

### Authors’ contributions

ASK analyzed and visualized the data. ASK and TS wrote the original draft. TS conceptualized the study. CH processed the raw data. CH and SL contributed to methodological discussions. TS, ML, and SL supervised the project. All authors contributed to supervision, review, and editing of the manuscript.

## Acknowledgements

We thank the Staatliches Amt für Landwirtschaft und Umwelt Mittleres Mecklenburg for kindly providing the inflow volume data.

