## Supplemental Information for "Enhancing the Understanding of Environmental Microbiomes through Topic Modeling: A Quantitative and Qualitative Analysis"

**Table 1.** Random Forest Performances when predicting several target variables with the full microbiome (*none*, *fractions*, *clr*).

| **Target** | $R^2$/Accuracy (***none***) | $R^2$/Accuracy (***fractions***) | $R^2$/Accuracy (***clr***) |
|---|---|---|---|
| Salinity | 0.91 | 0.92 | 0.89 |
| Chlorophyll a [mg/m$^3$] | 0.57 | 0.49 | 0.57 |
| Temperature [°C] | -1.3 | -0.9 | -0.6 |
| Phosphate [$\mu$mol/l] | 0.01 | -0.00 | -0.00 |
| Ammonium [$\mu$mol/l] | -0.34 | -0.89 | -0.54 |
| Nitrate [$\mu$mol/l] | 0.09 | 0.12 | -0.03 |
| Nitrite [$\mu$mol/l] | 0 | 0 | 0 |
| Location ID | 0.41 | 0.38 | 0.34 |
| Calendar week | 0 | 0 | 0 |

Table 2: Performance metrics ($R^2$, MAE, RSME) when predicting chlorophyll a with Random Forest based on the DRM approaches across $k \in \{2, 3, 4, \ldots, 20\}$.

| $k$ | Preprocession | Topic Model | Clustering Method | $R^2$ | Mean Absolute error | Root Mean Squared Error |
|---|---|---|---|---|---|---|
| 2 | fractions | none | pcoa | 0.4691931291545056 | 2.2271461187214605 | 4.603642864685615 |
| 2 | none | none | pca | 0.1972605096559211 | 2.8921004566210047 | 5.661358687786954 |
| 2 | fractions | none | pca | 0.0734250963868822 | 3.089383561643835 | 6.082381041370237 |
| 2 | clr | none | pca | -0.2781110580545907 | 3.817762557077626 | 7.143608632353639 |
| 2 | clr | lda | none | -2.982376686515864 | 6.095844748858448 | 12.60969336132591 |
| 2 | none | nnmf | none | 0.1212428851519551 | 2.899246575342465 | 5.923355309945022 |
| 2 | fractions | nnmf | none | 0.2392761231558812 | 2.679634703196347 | 5.511209001438273 |
| 2 | clr | nnmf | none | -1.098528008124541 | 5.274771689497718 | 9.153574955987734 |
| 2 | none | lda | none | -0.7769388591479114 | 3.975568130753062 | 8.423054285578184 |
| 2 | fractions | lda | none | -2.8462401774677937 | 5.658219178082192 | 12.392289693015428 |
| 3 | clr | lda | none | -0.0554793848785781 | 2.956164383561644 | 6.49169690967803 |
| 3 | fractions | none | pcoa | 0.2846597037033557 | 2.935730593607306 | 5.344286134951743 |
| 3 | none | none | pca | 0.2422896648666093 | 2.805593607305936 | 5.500282080769725 |
| 3 | fractions | none | pca | 0.1847435290366953 | 2.9018949771689497 | 5.705326258584601 |
| 3 | clr | none | pca | 0.3346670361562431 | 2.804269406392694 | 5.154100442352963 |
| 3 | none | nnmf | none | 0.2783217847670344 | 2.6560273972602744 | 5.367909127567873 |
| 3 | fractions | nnmf | none | 0.349688433025804 | 2.5864155251141554 | 5.09558554134188 |
| 3 | clr | nnmf | none | 0.0608201241397243 | 3.2902739726027392 | 6.123613140911449 |
| 3 | none | lda | none | 0.3434621104379826 | 2.538904109589041 | 5.119920937959775 |
| 3 | fractions | lda | none | -0.9164920341587904 | 5.746164383561643 | 8.747558720102646 |
| 4 | fractions | none | pcoa | 0.4451426178849561 | 2.383401826484018 | 4.706781540049785 |
| 4 | none | none | pca | 0.3003144538807762 | 2.7277625570776256 | 5.285484559807499 |
| 4 | fractions | none | pca | 0.27614841786083 | 2.791438356164384 | 5.375985902997119 |
| 4 | clr | none | pca | 0.2152290027776342 | 3.017602739726028 | 5.59763826176621 |
| 4 | fractions | nnmf | none | -0.1741629420832728 | 3.254954337899544 | 6.846956078013542 |
| 4 | clr | lda | none | 0.5019643952888317 | 2.3397945205479456 | 4.459267784971226 |
| 4 | fractions | lda | none | -1.085673676707195 | 6.19607305936073 | 9.125497221706825 |
| 4 | clr | nnmf | none | 0.1208378880737192 | 3.0798858447488584 | 5.924720115540361 |
| 4 | none | nnmf | none | 0.482605791124417 | 2.3528995433789954 | 4.545107284576515 |
| 4 | none | lda | none | 0.4229731478130636 | 2.385981735159817 | 4.799890932931344 |
| 5 | none | none | pca | 0.3335034629562613 | 2.662716894977169 | 5.15860536974196 |
| 5 | clr | lda | none | 0.4761515011444227 | 2.2936757990867584 | 4.573368636457323 |
| 5 | fractions | lda | none | -1.1438165979699615 | 6.0492694063926935 | 9.25181993697017 |
| 5 | none | lda | none | 0.5415363197992207 | 1.9993150684931509 | 4.278443738313945 |
| 5 | clr | nnmf | none | 0.4697451639712805 | 2.213767123287671 | 4.601248366441477 |
| 5 | none | nnmf | none | 0.4415333142987845 | 2.372716894977169 | 4.7220653470226885 |
| 5 | clr | none | pca | 0.2999605660545845 | 2.9154794520547944 | 5.2868210403160765 |
| 5 | fractions | none | pca | 0.2464849236189769 | 2.7867808219178083 | 5.48503407997138 |
| 5 | fractions | nnmf | none | 0.238687165935834 | 2.700296803652968 | 5.513341995046009 |
| 5 | fractions | none | pcoa | 0.4708040998694996 | 2.592328767123288 | 4.596651650135571 |
| 6 | fractions | lda | none | -0.5568351525035524 | 5.574155251141552 | 7.88414625057505 |
| 6 | clr | lda | none | 0.3909665934267358 | 2.6017123287671238 | 4.931214735450413 |
| 6 | fractions | none | pcoa | -0.3977033569737731 | 4.214657534246576 | 7.470348478665106 |

| | | | | | | |
|---|---|---|---|---|---|---|
| 6 | none | none | pca | 0.3181738666610268 | 2.683264840182648 | 5.21759275920546 |
| 6 | fractions | none | pca | 0.1883587223666132 | 2.919018264840183 | 5.692662284140746 |
| 6 | clr | none | pca | 0.2625511381563589 | 2.7057762557077623 | 5.426243918159295 |
| 6 | none | nnmf | none | -0.0930974984059544 | 3.213607305936073 | 6.606368692855105 |
| 6 | fractions | nnmf | none | 0.3578816212422432 | 2.6490867579908675 | 5.063384496271009 |
| 6 | clr | nnmf | none | 0.4994308047932597 | 2.4124657534246574 | 4.470595917087707 |
| 6 | none | lda | none | 0.4757611312161901 | 2.5506164383561645 | 4.575072347657616 |
| 7 | fractions | none | pcoa | 0.3543790821274202 | 2.302716894977169 | 5.077175242936626 |
| 7 | fractions | none | pca | 0.1373584992002358 | 3.028584474885845 | 5.868789479470012 |
| 7 | clr | none | pca | 0.1259533935545728 | 3.115616438356164 | 5.90745813729015 |
| 7 | none | none | pca | 0.3406349419485424 | 2.586575342465753 | 5.13093273940087 |
| 7 | none | nnmf | none | 0.4082303015135208 | 2.5754109589041096 | 4.860822012712398 |
| 7 | clr | lda | none | 0.3015783128339856 | 2.825388127853881 | 5.280708752762914 |
| 7 | fractions | lda | none | -1.2094043726197028 | 7.026666666666667 | 9.392278500650946 |
| 7 | clr | nnmf | none | 0.3919876634190437 | 2.6681278538812787 | 4.927079307488677 |
| 7 | fractions | nnmf | none | 0.2258432645112978 | 3.29662100456621 | 5.559654525911531 |
| 7 | none | lda | none | 0.5728053295953552 | 2.117694063926941 | 4.12996403955731 |
| 8 | clr | lda | none | 0.5818283998602567 | 2.281187214611872 | 4.086115369490366 |
| 8 | fractions | lda | none | -4.303631581371521 | 9.36849315068493 | 14.551911045237087 |
| 8 | none | lda | none | 0.2727075111403025 | 2.870057077625571 | 5.388748418958808 |
| 8 | clr | nnmf | none | 0.5994298189916233 | 2.174086757990868 | 3.999195781026126 |
| 8 | none | nnmf | none | 0.4811884500237343 | 2.3036529680365296 | 4.551328422665069 |
| 8 | clr | none | pca | 0.2400947746638404 | 3.034657534246575 | 5.5082427648175045 |
| 8 | fractions | none | pca | 0.2401844074145446 | 2.62175799086758 | 5.50791789962787 |
| 8 | none | none | pca | 0.3139016981480872 | 2.781392694063927 | 5.233913362557503 |
| 8 | fractions | none | pcoa | 0.4442054109817597 | 2.3615525114155247 | 4.710754963778856 |
| 8 | fractions | nnmf | none | 0.4708301748515227 | 2.4340182648401822 | 4.5965384037124775 |
| 9 | fractions | none | pcoa | 0.4921324459160318 | 2.220616438356165 | 4.503068892661402 |
| 9 | fractions | lda | none | 0.0199075967432905 | 3.471780821917808 | 6.25556971917151 |
| 9 | none | lda | none | 0.4636630829232679 | 2.649429223744292 | 4.627561539200398 |
| 9 | clr | nnmf | none | 0.4808657263756182 | 2.5508219178082188 | 4.552743766025798 |
| 9 | fractions | nnmf | none | 0.4748777096816139 | 2.401963470319634 | 4.578925568376341 |
| 9 | none | nnmf | none | 0.5439971691004256 | 2.126849315068493 | 4.266945802798328 |
| 9 | clr | none | pca | 0.0567694443008365 | 3.346392694063927 | 6.136804494718135 |
| 9 | fractions | none | pca | 0.3004521823932772 | 2.5036529680365294 | 5.28496432771362 |
| 9 | clr | lda | none | 0.4191296325627082 | 2.6607534246575346 | 4.815850185969102 |
| 9 | none | none | pca | -0.1760650033760866 | 3.582922374429224 | 6.852499627256649 |
| 10 | clr | lda | none | 0.3326359837571848 | 2.8231963470319634 | 5.16196137194763 |
| 10 | clr | none | pca | -0.0297364986299824 | 3.569794520547945 | 6.412042768553047 |
| 10 | fractions | lda | none | -0.6861013733012764 | 4.792406098515691 | 8.204936008928922 |
| 10 | none | lda | none | 0.4718056879618619 | 2.558219178082192 | 4.592299639803043 |
| 10 | clr | nnmf | none | 0.4264402970016276 | 2.650502283105023 | 4.785448786824931 |
| 10 | fractions | nnmf | none | 0.4489305694071717 | 2.3533561643835617 | 4.690687680206604 |
| 10 | none | nnmf | none | 0.516962530529693 | 2.2025342465753424 | 4.3916100219025465 |
| 10 | fractions | none | pca | 0.3777528789951011 | 2.512077625570776 | 4.984422011165199 |
| 10 | none | none | pca | 0.1082084419399259 | 3.295296803652968 | 5.967123627306681 |
| 10 | fractions | none | pcoa | 0.5457222464527591 | 2.22689497716895 | 4.258867139870472 |
| 11 | none | lda | none | 0.4512989668861514 | 2.7150000000000003 | 4.680596959741714 |
| 11 | fractions | none | pca | 0.3702481846680232 | 2.507853881278539 | 5.014389568005864 |
| 11 | clr | none | pca | -0.236858789653352 | 3.7262100456621 | 7.027379656755614 |
| 11 | none | nnmf | none | 0.5764206762983051 | 2.1513470319634704 | 4.126988492625687 |
| 11 | clr | lda | none | 0.5571835734193099 | 2.313744292237443 | 4.20479879146581 |
| 11 | fractions | nnmf | none | 0.4656064353224828 | 2.4683333333333337 | 4.619170224363229 |
| 11 | clr | nnmf | none | 0.460596785119526 | 2.479223744292238 | 4.6407708283515285 |
| 11 | fractions | lda | none | 0.2080416405815083 | 3.043006052116385 | 5.623212955501695 |
| 11 | none | none | pca | 0.0489085748684423 | 3.2184246575342463 | 6.16232345510172 |
| 11 | fractions | none | pcoa | 0.0829611976412825 | 3.1595205479452053 | 6.051000841779367 |
| 12 | clr | none | pca | 0.0630462414207448 | 3.386301369863013 | 6.116351502960124 |
| 12 | fractions | none | pcoa | 0.3867215177350297 | 2.802899543378995 | 4.9483706326232495 |
| 12 | fractions | none | pca | 0.3582987523362363 | 2.5420547945205483 | 5.061739598453982 |

| | | | | | | |
|---|---|---|---|---|---|---|
| 12 | none | nnmf | none | 0.576483480283184 | 2.127511415525114 | 4.112146086144065 |
| 12 | clr | nnmf | none | 0.4308173944444182 | 2.9152511415525115 | 4.7671538372243205 |
| 12 | fractions | nnmf | none | 0.2396072182455942 | 2.88324200913242 | 5.510009530194212 |
| 12 | clr | lda | none | 0.3604067888324379 | 2.865388127853881 | 5.05341866291013 |
| 12 | none | none | pca | 0.1219995571138297 | 3.2071232876712323 | 5.920804547020469 |
| 12 | fractions | lda | none | -3.322821056436248 | 8.782428127491483 | 13.137628785687571 |
| 12 | none | lda | none | 0.4341688873640361 | 2.8856164383561644 | 4.753098004382974 |
| 13 | fractions | none | pcoa | 0.3330901230274176 | 3.258675799086758 | 5.160204723205091 |
| 13 | none | none | pca | -0.1042501913637978 | 3.385570776255708 | 6.639985008510336 |
| 13 | clr | none | pca | 0.1298138861342318 | 3.2226027397260277 | 5.894397657461892 |
| 13 | clr | lda | none | 0.4908307896752847 | 2.480616438356164 | 4.508835845862157 |
| 13 | fractions | none | pca | 0.4007711183228291 | 2.518287671232877 | 4.891361106105297 |
| 13 | none | nnmf | none | 0.4523048950906529 | 2.385890410958904 | 4.6763045453555785 |
| 13 | fractions | nnmf | none | 0.3451865645067544 | 2.6856392694063924 | 5.113192557282963 |
| 13 | clr | nnmf | none | 0.4449381408493952 | 2.7060730593607305 | 4.707648735940753 |
| 13 | none | lda | none | 0.3601022318987706 | 2.9539041095890406 | 5.054621670146095 |
| 13 | fractions | lda | none | -0.2225880377651716 | 3.973173515981735 | 6.986721440276424 |
| 14 | fractions | none | pca | 0.4296505332806944 | 2.342716894977169 | 4.772037822269324 |
| 14 | none | none | pca | -0.2240653970700892 | 3.645022831050228 | 6.990941496944358 |
| 14 | none | lda | none | 0.2528084841605919 | 2.932214611872146 | 5.46197016071396 |
| 14 | clr | none | pca | -0.1307342017944175 | 3.5987899543378994 | 6.719138939045076 |
| 14 | fractions | none | pcoa | 0.3017050852794788 | 2.9521004566210047 | 5.280229473012629 |
| 14 | fractions | lda | none | -0.3251053047878516 | 3.9982891117961934 | 7.273753070559401 |
| 14 | none | nnmf | none | 0.4003425165240251 | 2.339931506849315 | 4.893110080072566 |
| 14 | fractions | nnmf | none | 0.3561633371089105 | 2.5802283105022834 | 5.070154681383072 |
| 14 | clr | nnmf | none | 0.2292752296287243 | 3.28148401826484 | 5.547317402609734 |
| 14 | clr | lda | none | 0.496056216072679 | 2.396141552511416 | 4.4856398728044535 |
| 15 | clr | lda | none | 0.426304386876031 | 2.991735159817352 | 4.786015730787192 |
| 15 | fractions | lda | none | -2.3588818223373598 | 8.018607704320035 | 11.580590884562222 |
| 15 | none | lda | none | 0.2739634144220781 | 3.192625570776256 | 5.3840937091114185 |
| 15 | clr | nnmf | none | 0.4136951717260967 | 2.8781050228310505 | 4.838325614182912 |
| 15 | fractions | nnmf | none | 0.5586804718061515 | 2.108493150684932 | 4.1976858160235935 |
| 15 | none | nnmf | none | 0.3623647340724691 | 2.385296803652968 | 5.0456778827828135 |
| 15 | clr | none | pca | -0.170006138211191 | 3.747260273972602 | 6.834825441281247 |
| 15 | fractions | none | pca | 0.1114723835328435 | 3.232579908675799 | 5.956193833075696 |
| 15 | none | none | pca | -0.0264594051820974 | 3.741552511415525 | 6.401831608626197 |
| 15 | fractions | none | pcoa | 0.4000082020889437 | 2.630684931506849 | 4.894473866435875 |
| 16 | fractions | none | pcoa | 0.1664408301256591 | 3.784520547945205 | 5.769013748874516 |
| 16 | clr | none | pca | -0.6412927454869317 | 4.4527168949771685 | 8.09517761910951 |
| 16 | fractions | none | pca | 0.3842076378673158 | 2.5032191780821917 | 4.958502153605052 |
| 16 | fractions | nnmf | none | 0.4753257876621657 | 2.210867579908676 | 4.5769715915408185 |
| 16 | none | nnmf | none | 0.3815400128150297 | 2.374018264840183 | 4.9692307132270015 |
| 16 | clr | lda | none | 0.5362891528137449 | 2.4968036529680364 | 4.302857710276491 |
| 16 | none | none | pca | -0.0374716935108638 | 3.605022831050229 | 6.436080765220272 |
| 16 | none | lda | none | 0.3073485405279228 | 3.0387442922374426 | 5.258849401909458 |
| 16 | clr | nnmf | none | 0.3356200081308074 | 3.0743607305936065 | 5.150407950327286 |
| 16 | fractions | lda | none | 0.1413004932533099 | 2.9986073059360727 | 5.855364885744139 |
| 17 | clr | lda | none | 0.5755083445739003 | 2.116872146118721 | 4.116877418671255 |
| 17 | fractions | lda | none | -1.4276757630474737 | 5.889987214192476 | 9.845294202597776 |
| 17 | none | lda | none | 0.326645245201022 | 2.8947945205479453 | 5.185078344916533 |
| 17 | clr | nnmf | none | 0.2741963795191385 | 3.4685844748858443 | 5.383229836289701 |
| 17 | none | nnmf | none | 0.4003544789938478 | 2.347785388127854 | 4.893061273899703 |
| 17 | clr | none | pca | -0.2343159871411049 | 3.965799086757991 | 7.020152302917964 |
| 17 | fractions | none | pca | 0.3360837940007488 | 2.682739726027397 | 5.148609955598292 |
| 17 | none | none | pca | -0.0567177177967548 | 3.76351598173516 | 6.4955039596245125 |
| 17 | fractions | none | pcoa | 0.1657378177320531 | 2.6177853881278543 | 5.771445989765943 |
| 17 | fractions | nnmf | none | 0.3928923394948872 | 2.268173515981735 | 4.923412383471132 |
| 18 | fractions | lda | none | 0.0195010235754313 | 3.6053164090590943 | 6.256867088214808 |
| 18 | fractions | none | pca | 0.3120092491457951 | 2.7847260273972605 | 5.24112668269997 |
| 18 | fractions | none | pcoa | 0.3178881340345566 | 2.697990867579908 | 5.218685912028222 |

| | | | | | | |
|---|---|---|---|---|---|---|
| 18 | none | none | pca | -0.391337264476832 | 4.1901826484018265 | 7.453316536768064 |
| 18 | clr | none | pca | -0.1597550620933618 | 4.141849315068493 | 6.804817709823456 |
| 18 | clr | lda | none | 0.4245700803524289 | 2.580136986301369 | 4.793244437481039 |
| 18 | none | nnmf | none | 0.3501717980050539 | 2.4451598173515983 | 5.09369146043063 |
| 18 | fractions | nnmf | none | 0.3941521042184635 | 2.31703196347032 | 4.918301624298769 |
| 18 | clr | nnmf | none | 0.3435655640727981 | 2.961735159817352 | 5.119517537556796 |
| 18 | none | lda | none | 0.3876446750262019 | 2.819703196347032 | 4.944644882466144 |
| 19 | none | nnmf | none | 0.3777810182465343 | 2.358127853881278 | 4.984309307156448 |
| 19 | clr | lda | none | 0.5907415434904193 | 2.1455022831050226 | 4.0423339419981295 |
| 19 | fractions | lda | none | -1.5564351521669368 | 5.601565459219914 | 10.103009179241226 |
| 19 | none | lda | none | 0.4015390208252064 | 2.73337899543379 | 4.888225999752623 |
| 19 | clr | nnmf | none | 0.3686941386122482 | 2.95351598173516 | 5.020572790185649 |
| 19 | fractions | nnmf | none | 0.4026637045618685 | 2.2204337899543383 | 4.883630634427218 |
| 19 | clr | none | pca | -0.5660826162869943 | 4.461347031963471 | 7.907527149269762 |
| 19 | fractions | none | pca | 0.3995387266121792 | 2.461735159817352 | 4.896388381050077 |
| 19 | none | none | pca | -0.1096118709786577 | 3.785707762557078 | 6.656085689977914 |
| 19 | fractions | none | pcoa | 0.3046439987905619 | 2.9468493150684933 | 5.269106307386369 |
| 20 | none | none | pca | -0.2900033427174446 | 3.874931506849314 | 7.176765816153449 |
| 20 | fractions | none | pca | 0.3955038232810549 | 2.570525114155251 | 4.912811901167773 |
| 20 | clr | none | pca | -0.1916914942199203 | 4.12513698630137 | 6.897874316869234 |
| 20 | fractions | nnmf | none | 0.3914039154772031 | 2.3984018264840183 | 4.929443965420358 |
| 20 | none | nnmf | none | 0.3767087589474128 | 2.462762557077625 | 4.988602145878445 |
| 20 | clr | nnmf | none | 0.3793822051250718 | 2.8453424657534248 | 4.977891991174486 |
| 20 | none | lda | none | 0.3750616949678292 | 2.9601826484018265 | 4.995189055344166 |
| 20 | fractions | lda | none | 0.581546938727552 | 2.2237899543378994 | 4.087490270609677 |
| 20 | clr | lda | none | 0.4624562744296294 | 2.447579908675799 | 4.6327648381568105 |
| 20 | fractions | none | pcoa | 0.1428786758220497 | 3.378812785388128 | 5.849981696103746 |

**Table 3.** The approaches, where $R^2 > baseline$, their bootstrapped mean $R^2$, t-test p-values, and significance at $\alpha = 0.05$. The test evaluates whether the bootstrapped $R^2$ distributions differ from the full microbiome baseline.

| **Approach** | $R^2$ | Bootstrapped mean $R^2$ | p-value | significant |
|---|---|---|---|---|
| *clr* + nnmf + $k$=8 | 0.6 | 0.57 | 0.02 | yes |
| *none* + nnmf + $k$=12 | 0.58 | 0.51 | $< 0.01$ | yes |
| *clr* + lda + $k$=19 | 0.59 | 0.55 | $< 0.01$ | yes |
| *fractions* + lda + $k$=20 | 0.58 | 0.59 | $< 0.01$ | yes |

**Table 4.** Bacterial identification numbers of the top 5 contributing ASVs across extracted topics.

| Phylum | Class | Ordner | Family | Genus | ID |
|---|---|---|---|---|---|
| Proteobacteria | Alphaproteobacteria | Rhodobacterales | Rhodobacteraceae | *Pseudorhodobacter* | 9129c25a-b415-46d9-8bce-618c0704cbfd |
| Proteobacteria | Alphaproteobacteria | Rhodobacterales | Rhodobacteraceae | *Pseudorhodobacter* | 7fb18c69-c34c-4630-94ef-c4515c4066c8 |
| Proteobacteria | Alphaproteobacteria | Rhodobacterales | Rhodobacteraceae | *Tabrizicola* | da897aba-58df-408a-a5b6-023b6da4b80c |
| Proteobacteria | Alphaproteobacteria | Rhodobacterales | Rhodobacteraceae | *Yoonia-Loktanella* | e6368140-273a-40c7-a909-d8dd48618e79 |
| Proteobacteria | Alphaproteobacteria | Rhodobacterales | Rhodobacteraceae | *Yoonia-Loktanella* | 6c8700a1-b831-47ad-a78d-91c850bf6459 |
| Proteobacteria | Alphaproteobacteria | Rhodobacterales | Rhodobacteraceae | *Planktomarina* | 307a04d-0ecb-45fd-95e3-519fc0d1ba32 |
| Proteobacteria | Alphaproteobacteria | Rhodobacterales | Rhodobacteraceae | *Planktomarina* | f202e1c3-12fe-4a09-904d-86cd64146552 |
| Proteobacteria | Alphaproteobacteria | Rhodobacterales | Rhodobacteraceae | *Lentibacter* | 90fdc862-54ed-4ed3-b394-8314995dc79a |
| Proteobacteria | Alphaproteobacteria | Rhodobacterales | Rhodobacteraceae | *Seohaeicola* | 4e7a02b2-0b3e-4f26-b715-5d0238df4b47 |
| Proteobacteria | Alphaproteobacteria | Rhodobacterales | Rhodobacteraceae | *Marivita* | c5055337-5e3f-409a-a65c-52ee5ee92744 |
| Proteobacteria | Alphaproteobacteria | Rhodobacterales | Rhodobacteraceae | *Rhodobacter* | 05fab431-d352-4c80-954e-0107499d1b6c |
| Proteobacteria | Gammaproteobacteria | Burkholderiales | Burkholderiaceae | *Polynucleobacter* | fcecd815-81d7-4dda-8a51-761486ec0e05 |
| Proteobacteria | Gammaproteobacteria | Burkholderiales | Comamonadaceae | *RS62 marine group* | ae897dbb-82f8-49ce-a62e-592e687bbb5f |
| Proteobacteria | Gammaproteobacteria | Pseudomonadales | SAR86 clade | SAR86 clade (unclassified) | 6715e23a-6d57-446b-90a7-0f4359c6a3cd |
| Cyanobacteria | Cyanobacteriia | Synechococcales | Cyanobiaceae | *Cyanobium PCC-6307* | 167cd9db-5f68-4258-b1a8-d50ba42d3f6f |
| Cyanobacteria | Cyanobacteriia | Synechococcales | Cyanobiaceae | *Cyanobium PCC-6307* | d4be031f-3dde-45ae-aadc-9e4b111a7d17 |
| Cyanobacteria | Cyanobacteriia | Synechococcales | Cyanobiaceae | *Cyanobium PCC-6307* | c4e5479a-310f-43eb-8569-9a5df2af6873 |
| Cyanobacteria | Cyanobacteriia | Synechococcales | Cyanobiaceae | *Cyanobium PCC-6307* | d5971909-dd7b-40f5-8351-6a992a9e25a2 |
| Cyanobacteria | Cyanobacteriia | Synechococcales | Cyanobiaceae | *Synechococcus CC9902* | 31c638bb-18d2-4f9c-a320-d454c56aa9b9 |
| Bacteroidota | Bacteroidia | Flavobacteriales | Flavobacteriaceae | *NS3a marine group* | 9fbd4c78-3fce-4db9-a574-c0b8b005040e |
| Bacteroidota | Bacteroidia | Flavobacteriales | Flavobacteriaceae | *NS3a marine group* | 1d8af072-a47a-44dd-9a65-ac125b94bdec |
| Bacteroidota | Bacteroidia | Flavobacteriales | Flavobacteriaceae | *NS5 marine group* | 1a86ec6b-4ce3-40ea-bd95-f5daf3c1d87d |
| Bacteroidota | Bacteroidia | Flavobacteriales | Flavobacteriaceae | *Polaribacter* | fd77a141-bf9a-43ce-89ea-fe48914b4eca |
| Bacteroidota | Bacteroidia | Flavobacteriales | Flavobacteriaceae | Flavobacteriaceae (unclassified) | d75b3c67-0803-4297-bc7c-c6a863521fa2 |
| Bacteroidota | Bacteroidia | Flavobacteriales | Cryomorphaceae | Cryomorphaceae (unclassified) | 865ea2ae-30d7-4a54-ad63-3c5b2ac3636c |
| Bacteroidota | Bacteroidia | Flavobacteriales | Crocinitomicaceae | Crocinitomicaceae (unclassified) | a92c9174-9b90-4da4-842b-20ff64afc329 |
| Actinobacteriota | Acidimicrobiia | Microtrichales | Ilumatobacteraceae | *Ilumatobacter* | 6a2396a5-2e01-4512-9981-0b79f530c193 |
| Actinobacteriota | Acidimicrobiia | Microtrichales | Ilumatobacteraceae | Ilumatobacteraceae (unclassified) | 49ad3683-413f-4e97-b61d-2536d67b98a9 |
| Actinobacteriota | Actinobacteria | Corynebacteriales | Mycobacteriaceae | *Mycobacterium* | adbd1e81-ffce-460e-89e8-5ecf19c6bb65 |
| Actinobacteriota | Actinobacteria | Corynebacteriales | Mycobacteriaceae | *Mycobacterium* | a6e3477b-3b31-43a5-9711-bda88ee53f66 |
| Actinobacteriota | Actinobacteria | Micrococcales | Microbacteriaceae | *Candidatus* Limnoluna | 381985e7-57c3-410d-adec-0af27163e4db |
| Actinobacteriota | Actinobacteria | Frankiales | Sporichthyaceae | Sporichthyaceae (unclassified) | 9ef8efbe-4313-4521-856c-f9924d47c75f |
| Actinobacteriota | Actinobacteria | Micrococcales | Microbacteriaceae | *Candidatus* Aquiluna | da565a40-88bf-4491-8a27-cd67e928b714 |
| Actinobacteriota | Actinobacteria | Micrococcales | Microbacteriaceae | *Candidatus* Aquiluna | 6b70ca18-a760-4b45-954c-bc86f65afaea |
| Actinobacteriota | Actinobacteria | Micrococcales | Microbacteriaceae | *Candidatus* Aquiluna | 608fed8c-61ae-4e37-b740-d9c1c6f4d785 |
| Actinobacteriota | Acidimicrobiia | Microtrichales | Ilumatobacteraceae | *CL500-29 marine group* | d481a517-1f38-4b72-93ad-1c32e42bbbcd |
| Actinobacteriota | Acidimicrobiia | Microtrichales | Ilumatobacteraceae | *CL500-29 marine group* | 0e355dc3-7810-4603-b37a-03639af58466 |
| Actinobacteriota | Acidimicrobiia | Microtrichales | Ilumatobacteraceae | *CL500-29 marine group* | b7afcb6d-b3d5-4eb5-b346-6527f092b9fa |
| Actinobacteriota | Actinobacteria | Frankiales | Sporichthyaceae | *hgcI clade* | 06802502-e084-41f4-9a57-98a020f96856 |
| Actinobacteriota | Actinobacteria | Frankiales | Sporichthyaceae | *hgcI clade* | cc4e982f-16b2-419d-972f-b32e9c590c04 |
| Actinobacteriota | Actinobacteria | Frankiales | Sporichthyaceae | *hgcI clade* | db72cb19-c152-4378-8845-484741bc81ab |
| Actinobacteriota | Actinobacteria | PeM15 | PeM15 unclassified | PeM15 (unclassified) | 92c9e165-c9b3-44b4-8c15-1e6a25d411e5 |
| Actinobacteriota | Actinobacteria | PeM15 | PeM15 (unclassified) | PeM15 (unclassified) | e730717f-2fa8-444e-9dcc-c407e31b1d9b |
| Actinobacteriota | Actinobacteria | Micrococcales | Microbacteriaceae | *ML602J-51* | a5045aa2-f32c-4e0a-a8c4-c0f7f51a22e8 |
| Verrucomicrobiota | Verrucomicrobiae | Verrucomicrobiales | Rubritaleaceae | *Luteolibacter* | 623f6a46-892a-4217-9faf-5b372e1221b3 |
| Verrucomicrobiota | Verrucomicrobiae | Verrucomicrobiales | Rubritaleaceae | *Luteolibacter* | fbaaa28c-9fe5-401e-893d-98e6a3aee734 |
| Planctomycetota | Planctomycetes | Planctomycetales | Rubinisphaeraceae | *Fuerstia* | 8f15350d-04e5-448a-8267-5aff58f4d3a7 |

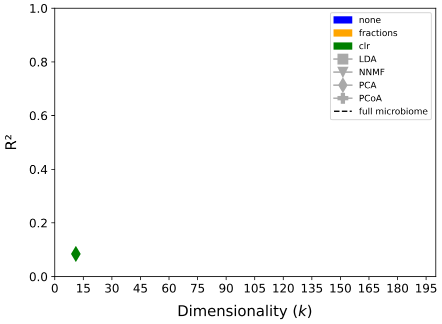


**Fig. 1.** RF performance ($R^2$) based on DMR generated topic or PCA/PCoA-component clusters across $k \in \{11, 21, \ldots, 191\}$ when predicting temperature. RF performance (Full microbiome)<0.15

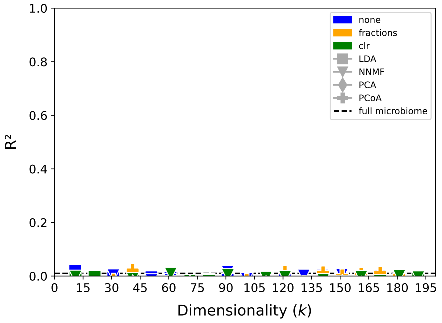


**Fig. 2.** RF performance ($R^2$) based on DMR generated topic or PCA/PCoA-component clusters across $k \in \{11, 21, \ldots, 191\}$ when predicting phosphate concentrations. RF performance (Full microbiome)<0.15

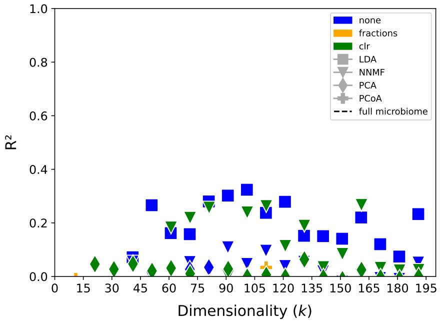


**Fig. 3.** RF performance ($R^2$) based on DMR generated topic or PCA/PCoA-component clusters across $k \in \{11, 21, \ldots, 191\}$ when predicting ammonium concentrations. RF performance (Full microbiome)<0.15

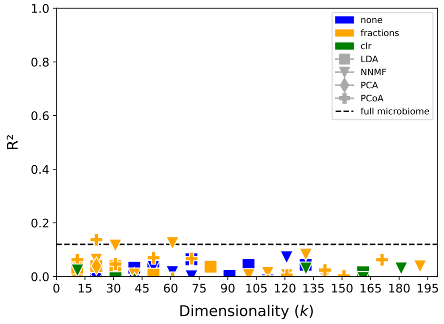


**Fig. 4.** RF performance ($R^2$) based on DMR generated topic or PCA/PCoA-component clusters across $k \in \{11, 21, \ldots, 191\}$ when predicting nitrate concentrations. RF performance (Full microbiome)<0.15

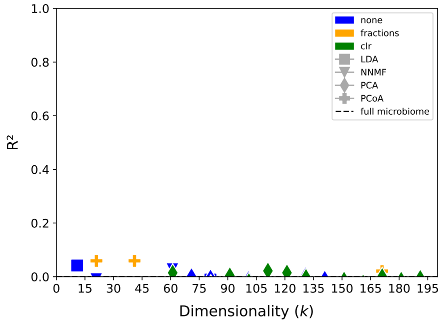


**Fig. 5.** RF performance ($R^2$) based on DMR generated topic or PCA/PCoA-component clusters across $k \in \{11, 21, \ldots, 191\}$ when predicting nitrite concentrations. RF performance (Full microbiome)<0.15

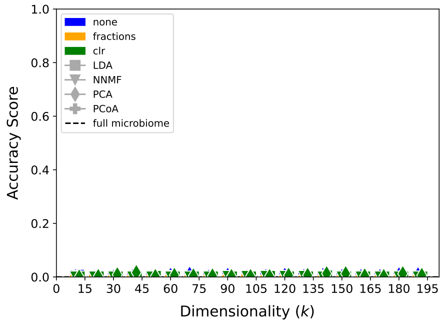


**Fig. 6.** RF performance (Accuracy) based on DMR generated topic or PCA/PCoA-component clusters across $k \in \{11, 21, \ldots, 191\}$ when predicting the calendar week. RF performance (Full microbiome)<0.1

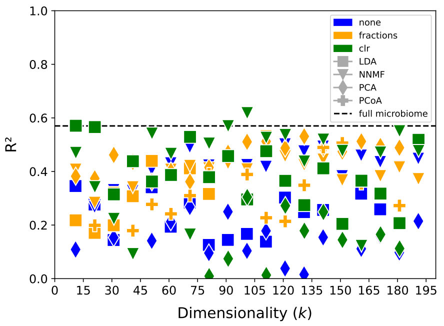


**Fig. 7.** RF performance ($R^2$) based on DMR generated topic or PCA/PCoA-component clusters across $k \in \{11, 21, \ldots, 191\}$ when predicting chlorophyll a.

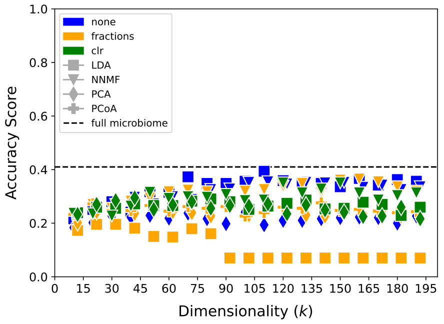


**Fig. 8.** RF performance (Accuracy) based on DMR generated topic or PCA/PCoA-component clusters across $k \in \{11, 21, \ldots, 191\}$ when predicting the location id.

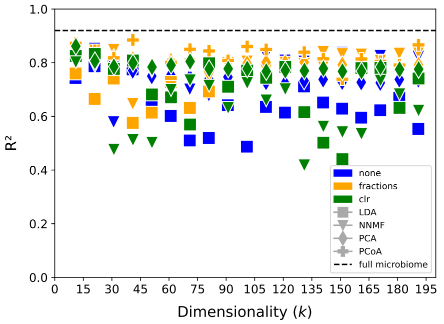


**Fig. 9.** RF performance ($R^2$) based on DMR generated topic or PCA/PCoA-component clusters across $k \in \{11, 21, \ldots, 191\}$ when predicting salinity.

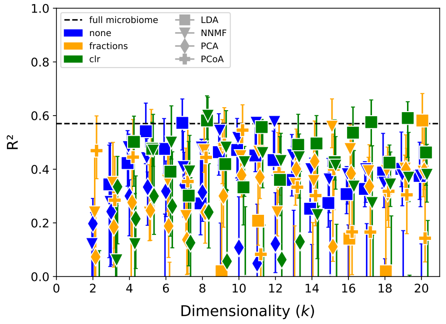


**Fig. 10.** RF performance ($R^2$) based on DMR generated topic or PCA/PCoA-component clusters across $k \in \{2, 3, 4, \ldots, 20\}$ when predicting chlorophyll a concentration, with 95% confidence intervals.

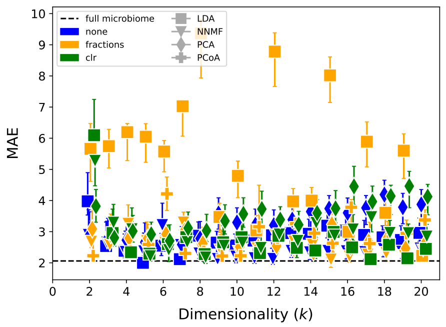


**Fig. 11.** RF performance (MAE) based on DMR generated topic or PCA/PCoA-component clusters across $k \in \{2, 3, 4, \ldots, 20\}$ when predicting chlorophyll a concentration, with 95% confidence intervals.

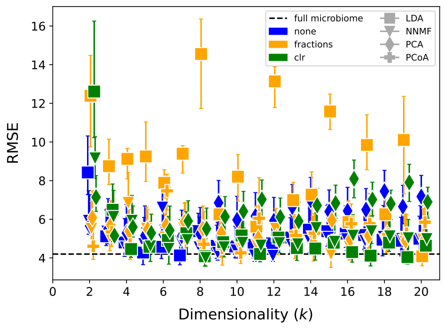


**Fig. 12.** RF performance (RMSE) based on DMR generated topic or PCA/PCoA-component clusters across $k \in \{2, 3, 4, \ldots, 20\}$ when predicting chlorophyll a concentration, with 95% confidence intervals.

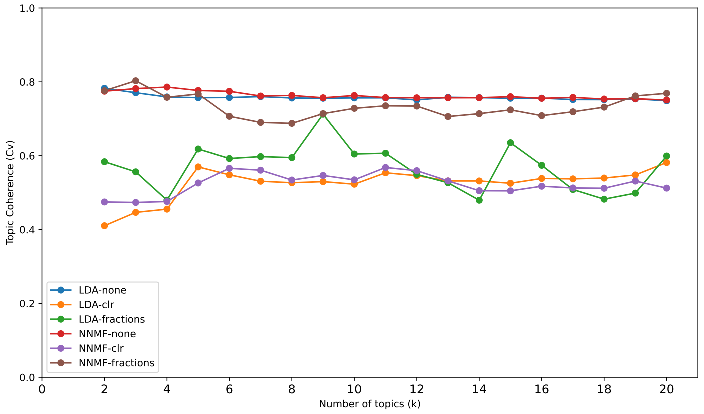


**Fig. 13.** The results of the topic coherence analysis using C_v as the coherence metric for LDA and NNMF topics based on various preprocessed microbiome data across $k \in \{2, 3, 4, \dots, 20\}$. The NNMF model with unprocessed data (red) selected in our study shows that topic coherence is among the highest performing approaches here. Furthermore the analysis suggests that the choice of k does not influence topic coherence drastically across the k range considered.
